# Molecular determinants of metazoan tricRNA biogenesis

**DOI:** 10.1101/497545

**Authors:** Casey A. Schmidt, Joseph D. Giusto, Alicia Bao, Anita K. Hopper, A. Gregory Matera

## Abstract

Mature tRNAs are generated by multiple post-transcriptional processing steps, which can include intron removal. Recently, we discovered a new class of circular non-coding RNAs in metazoans, called tRNA intronic circular (tric)RNAs. To investigate the mechanism of tricRNA biogenesis, we generated constructs that replace native introns of human and fruit fly tRNA genes with the Broccoli fluorescent RNA aptamer. Using these reporters, we identified *cis*-acting elements required for tricRNA formation *in vivo*. Disrupting a conserved base pair in the anticodon-intron helix dramatically reduces tricRNA levels. Although the integrity of this base pair is necessary for proper splicing, it is not sufficient. In contrast, strengthening weak bases in the helix also interferes with splicing and tricRNA production. Furthermore, we identified *trans*-acting factors important for tricRNA biogenesis, including several known tRNA processing enzymes such as the RtcB ligase and components of the TSEN endonuclease complex. Depletion of these factors inhibits *Drosophila* tRNA intron circularization. Notably, RtcB is missing from fungal genomes and these organisms normally produce linear tRNA introns. Here, we show that in the presence of exogenous RtcB, yeast lacking the tRNA ligase Rlg1/Trl1 are converted into producing tricRNAs. In summary, our work characterizes the major players in eukaryotic tricRNA biogenesis.

## INTRODUCTION

Accurate processing of RNAs is crucial for their proper function *in vivo*; most primary transcripts can be considered as precursor molecules containing sequences that must be removed. One example of this phenomenon is in the transcription and processing of tRNA genes. As the translators between the languages of nucleic acids and polypeptides, tRNAs are essential for protein expression. Thus, it is important that they are processed correctly. In eukaryotes, transcription of pre-tRNAs is carried out by RNA polymerase III (1). Following transcription, a pre-tRNA molecule contains 5′ leader and 3′ trailer sequences, which are removed by RNase P and RNase Z, respectively (2, 3). In some instances, the pre-tRNA transcript also contains an intron. Unlike messenger RNA splicing, which occurs by a large ribonucleoprotein complex called the spliceosome, tRNA splicing is carried out by a relatively small protein-only complex called TSEN. The pre-tRNA is first recognized by the TSEN (tRNA splicing endonuclease) complex, a heterotetramer consisting of two structural members, TSEN15 and TSEN54, and two catalytic members, TSEN2 and TSEN34 (4, 5). TSEN cleaves an intron-containing precursor into three segments: the 5′ exon, an intron, and the 3′ exon. Notably, this complex produces a 5′-OH and a 2′,3′-cyclic phosphate at each site of cleavage.

Ligation of the exon halves is handled differently in different organisms. Plants and fungi utilize the “healing and sealing” pathway, wherein a multifunctional enzyme called Rlg1/Trl1 performs three distinct activities: a cyclic phosphodiesterase to open the 2′,3′-cyclic phosphate into a 3′-OH and 2′-phosphate; a kinase to phosphorylate the 5′-OH of the intron and 3′ exon; and a ligase to join the exon halves together. A separate 2′-phosphotransferase enzyme called Tpt1 removes the remaining 2′-phosphate at the junction of the newly formed tRNA. The intron, now phosphorylated on its 5′ end, is degraded by a 5′ to 3′ exonuclease, creating a supply of nucleotides (6).

In contrast, archaea and animals are thought to use the “direct ligation” pathway, wherein a ligase enzyme directly joins the exon halves together using the cyclic phosphate as the junction phosphate. That is, there is no addition of an external phosphate onto the tRNA molecule. In archaea, the tRNA ligase is able to act on the intron ends to generate a circular RNA (7). Similarly, we have shown that RtcB, which ligates tRNA exon halves in *Drosophila*, also joins the intron ends together to make a circular RNA, which we term tRNA intronic circular (tric)RNA (8).

Although there have been many studies on the mechanism of pre-tRNA splicing in various organisms, much of this work comes from *in vitro* experiments, using purified proteins or cell extracts combined with an *in vitro* transcribed substrate. Thus, there is a need for an *in vivo* tRNA splicing model, where this processing pathway is placed in a cellular context. In this manuscript we present a unique system that allows us to detect both newly synthesized tRNAs and tricRNAs in *Drosophila* and human cultured cells. We take advantage of fluorescent RNA aptamer technology and northern blotting to elucidate the *cis*-acting elements and *trans*-acting factors necessary for proper tRNA and tricRNA biogenesis.

## MATERIALS AND METHODS

### Generation of reporter and mutant constructs

The human, heterologous, and tricRNA reporters used in this study were described previously (9). To generate the dual reporter, four point mutations were introduced in the 3′ exon of the tricRNA reporter (see Supplementary Figure S1) using Q5 Site-Directed Mutagenesis (NEB). For primer sequences, see Supplementary Table 1. Additional mutations within the heterologous, tricRNA, and dual reporters were also generated using Q5 Site-Directed Mutagenesis (NEB). See Supplementary Table 1 for primer sequences.

### Generation of *in vitro* transcribed RNAs

*In vitro*-transcribed RNAs were used for both S2 cell RNAi and analysis of pre-tRNA fluorescence. Templates were generated by PCR (see Supplementary Table 1 for primer sequences) and *in vitro* transcription was carried out using the MEGAscript T7 transcription kit (Invitrogen). RNA was isolated by phenol/chloroform extraction and ethanol precipitation.

### Cell culture and transfections

For human cell culture, HEK293T cells were maintained in DMEM (Gibco) supplemented with 10% fetal bovine serum (HyClone) and 1% penicillin/streptomycin (Gibco) at 37° and 5% CO_2_. Cells (2 × 10^6^) were plated in T25 flasks and transiently transfected with 2.5 µg plasmid DNA per flask using FuGENE HD transfection reagent (Promega) according to the manufacturer’s protocol. Cells were harvested 72 hours post-transfection. RNA was isolated using TRIzol Reagent (Invitrogen), with a second chloroform extraction and ethanol rather than isopropanol precipitation (9).

For *Drosophila* cell culture, S2 cells were maintained in SF-900 serum-free medium (Gibco) supplemented with 1% penicillin-streptomycin and filter sterilized. Cells (5 × 10^6^) were plated in T25 flasks and transiently transfected with 2.5 µg plasmid DNA per flask using Cellfectin II transfection reagent (Invitrogen) according to the manufacturer’s protocol. Cells were harvested 72 hours post-transfection. RNA was isolated using TRIzol Reagent (Invitrogen), with a second chloroform extraction and ethanol rather than isopropanol precipitation (9). S2 RNAi was performed as described in (10) for 10 days, with dsRNA targeting Gaussia luciferase used as a negative control. In experiments with both RNAi and reporter expression, the reporter was transfected on day 7, and cells were harvested on day 10. Primers used to make PCR products for *in vitro* transcription can be found in Supplementary Table 1.

### In-gel staining assay

RNA samples (5 µg) were electrophoresed through 10% TBE-urea gels (Invitrogen). Gels were washed 3X in dH_2_O to remove urea and then incubated in DFHBI-1T staining solution (40 mM HEPES pH 7.4, 100 mM KCl, 1 mM MgCl_2_, 10 µM DFHBI-1T (Lucerna)). Following staining, gels were imaged on an Amersham Typhoon 5. To visualize total RNA, gels were washed 3X in dH_2_O, stained with ethidium bromide, and imaged on an Amersham Imager 600. Gels were quantified using ImageQuant TL software (GE Healthcare). For analysis of the *in vitro* transcribed RNAs, the DFHBI-stained gel was subsequently stained in SYBR Gold (Invitrogen) to detect total RNA and imaged on an Amersham Imager 600.

### Northern blotting of *Drosophila* samples

RNA samples (5 µg) were separated by electrophoresis through 10% (for nuclease knockdown and overexpression experiments) or 15% (for dual reporter experiments) TBE-urea gels (Invitrogen). Following electrophoresis, the RNA was transferred to a nylon membrane (PerkinElmer). The membrane was dried overnight and UV-crosslinked. Pre-hybridization was carried out in Rapid-hyb Buffer (GE Healthcare) at 42°. Probes were generated by end-labeling oligonucleotides (IDT) with γ-^32^P ATP (PerkinElmer) using T4 PNK (NEB), and then probes were purified using Illustra Microspin G-50 columns (GE Healthcare) to remove unincorporated nucleotides. Upon purification, probes were boiled, cooled on ice, and then added to the rapid-hyb buffer for hybridization. After hybridization, the membrane was washed in SSC buffer. For probe sequences, see Supplementary Table 1. Washing conditions are as follows. U1 and U6: hybridization at 65°, washes (twice in 2X SSC, twice in 0.33X SSC) at 60°. 7SK and dual reporter probe: hybridization at 42°, two washes in 5X SSC at 25°, and two washes in 1X SSC at 42°. For the dual reporter probe, two additional washes in 0.1X SSC at 45° were performed. The membrane was then exposed to a storage phosphor screen (GE Healthcare) and imaged on an Amersham Typhoon 5.

### RT-PCR of *Drosophila* samples

To test knockdown efficiency, total RNA was treated with TURBO DNase (Invitrogen) and then converted to cDNA using the SuperScript III kit (Invitrogen) with random hexamer priming. Primers for each candidate processing factor can be found in Supplementary Table 1.

### Endonuclease overexpression cloning

To generate overexpression vectors, the Gateway system (Invitrogen) was used to clone ORFs of Dis3 and Clipper from S2 cell cDNA into either pAFW (*Drosophila* Genomics Resource Center, Barcode #1111) or pAWF (*Drosophila* Genomics Resource Center, Barcode #1112). Primers used to isolate Dis3 and Clipper can be found in Supplementary Table 1.

### Western blotting

To test overexpression of candidate endonucleases, cells were washed in ice-cold 1X PBS and collected in ice-cold 1X PBS by scraping. The harvested cells were split into two portions: one for RNA isolation (see above) and one for protein isolation. Cells were pelleted by spinning at 2000 RPM for 5 minutes. The supernatant was removed and cells were lysed in ice-cold lysis buffer (50 mM Tris-HCl pH 7.5, 150 mM NaCl, 1 mM EDTA, 1% NP-40, 2X protease inhibitor cocktail (Invitrogen)) for 30 minutes on ice. The lysate was cleared by centrifugation at 13,000 RPM for 10 minutes at 4°. Protein lysate samples were electrophoresed through 4-12% Bis-Tris gels (Invitrogen). Following electrophoresis, protein lysate samples were transferred to a nitrocellulose membrane (GE Healthcare). The membrane was blocked in 5% milk in TBST. Washes and antibody dilutions were performed in TBST. The following antibodies and dilutions were used in this study: anti-FLAG M2 at 1:10,000 (Sigma) and anti-β-tubulin at 1:20,000 (Sigma). Membranes were imaged on an Amersham Imager 600.

### Yeast strains

Wild type and *xrn1Δ S. cerevisiae* strains in the MATa/BY4741 background (MATa his3Δ1 leu2Δ0 met15Δ0 ura3Δ0) were purchased from Open Biosystems, then transformed with a multi-copy episomal plasmid with a His cassette as the selection marker. The *rlg1Δ+RtcB* strain was gifted by Dr. Stewart Shuman and used as described (11). Yeast strains were grown in synthetic define media (SC) lacking histidine.

### Northern blotting of yeast samples

Yeast mutant strains were grown in the 15 mL culture tubes to early log phase (0.4 OD600), and small RNAs were isolated using phenol extraction. Five micrograms of small RNAs were separated through electrophoresis on a 10% polyacrylamide gel. RNAs were transferred onto a Hybond N+ membrane. tRNA introns were detected using non-radioactive digoxygenin-labeled probes complementary to the full intron and half of the 5′ exon, using the method described in (12).

### Terminator Exonuclease (TEX) treatment

Five micrograms of small RNAs was dissolved in nuclease-free water and incubated with TEX (Lucigen) for 1 h at 30°C according to the manufacturer’s protocol. The reaction was terminated by adding 1 µL of 100 mM EDTA (pH 8.0)

### RT-PCR of yeast samples

Small RNAs were extracted from yeast cultures grown at 23 °C to an OD600 of 0.2-0.6. First-strand cDNA synthesis was carried out using 1 µg of RNA and RevertAid reverse transcriptase (Thermofisher), according to the manufacturer’s protocol. PCR was carried out using *Taq* DNA polymerase. 10 µM of the forward and reverse primer were added to 1 µL of cDNA sample. PCR conditions were as follows: 2 min at 95°C; 40 cycles of 30 sec at 95°C for denaturing, 30 sec at 55°C for annealing, and 30 s at 72°C for extension. Rolling circle concatemers were formed as a result of reverse transcriptase reading around the RNA multiple iterations during cDNA synthesis.

### Visualization of *Drosophila* BHB-like motifs

Sequences for all intron-containing pre-tRNAs were obtained from the genomic tRNA database (13, 14) and FlyBase (flybase.org). The structure of the sequence containing the anticodon loop and intron was predicted using Mfold (15) and then drawn using VARNA (16).

## RESULTS

### Generation and characterization of *in vivo* splicing reporters

To characterize the *cis*-acting elements and *trans*-acting factors necessary for proper tricRNA splicing, we first sought to develop a reporter system that would allow us to detect newly synthesized tRNAs and tricRNAs. Previously, we showed that human cells transfected with native fruit fly intronic tRNA genes readily express *Drosophila* tricRNAs (8). Moreover, when provided with either a human or fruit fly tRNA construct that contains a synthetic intron, human cells can also produce “designer” tricRNAs. For example, we inserted the 49 nt Broccoli fluorescent RNA aptamer (17) into the introns of the human (*TRYGTA3-1*) and the *Drosophila* (*CR31905*) tRNA:Tyr_GUA_ genes (9). We term the resultant circular RNA “tricBroccoli,” or tricBroc for short (18). This system is enhanced by use of external pol III promoters to increase overall expression levels (8, 9). Specifically, we used the human and *Drosophila* U6+27 promoters, which take advantage of a sequence element within the U6 snRNA that promotes 5′-methylphosphate capping of the transcript (19), enhancing the stability of the linear precursor to facilitate downstream processing events (9). A schematic of the reporters can be seen in Figure 2A. For more detailed sequence information, see Supplementary Figure S1.

**Figure 1:**
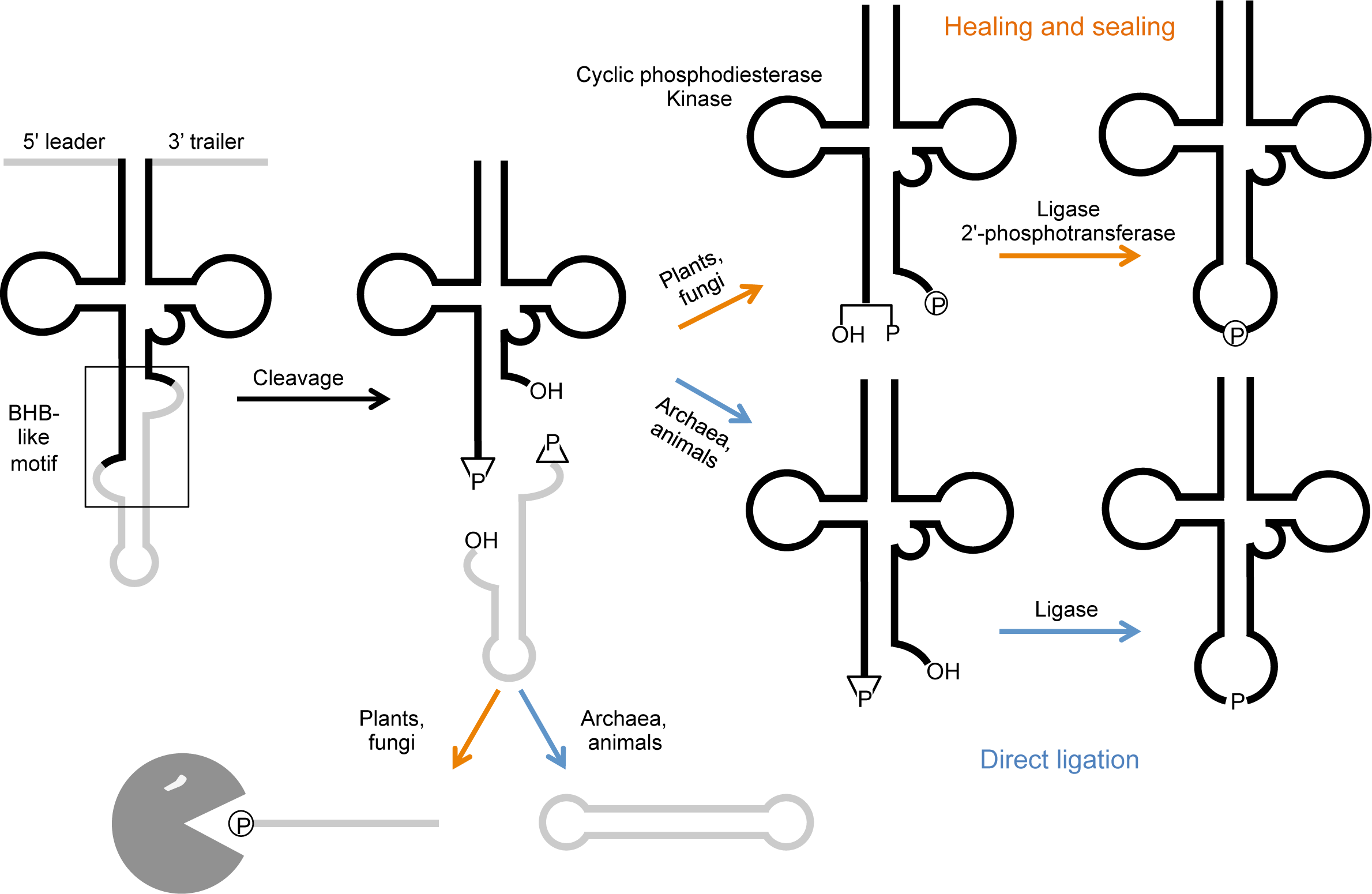
tRNA splicing pathway. Pre-tRNAs are transcribed by RNA polymerase III and contain a 5′ leader and 3′ trailer sequence. A structural motif resembling the archaeal bulge-helix-bulge (BHB) is present in the pre-tRNA. The leader and trailer are removed by RNase P and RNase Z, respectively. Cleavage of the pre-tRNA yields two exon halves and an intron, each bearing 5′-OH and 2′,3’-cyclic phosphate at the cut sites. In plants and fungi, a multifunctional enzyme phosphorylates the 5′ end of both the 3′ exon and intron, opens the cyclic phosphodiesterase, and ligates the exon halves to make a mature tRNA. A separate 2′-phosphotransferase removes the extra phosphate (“healing and sealing”). The intron is degraded by an exonuclease. In archaea and animals, a single enzyme joins both the exon halves and the intron ends to yield a mature tRNA and a circular RNA (“direct ligation”).

**Figure 2:**
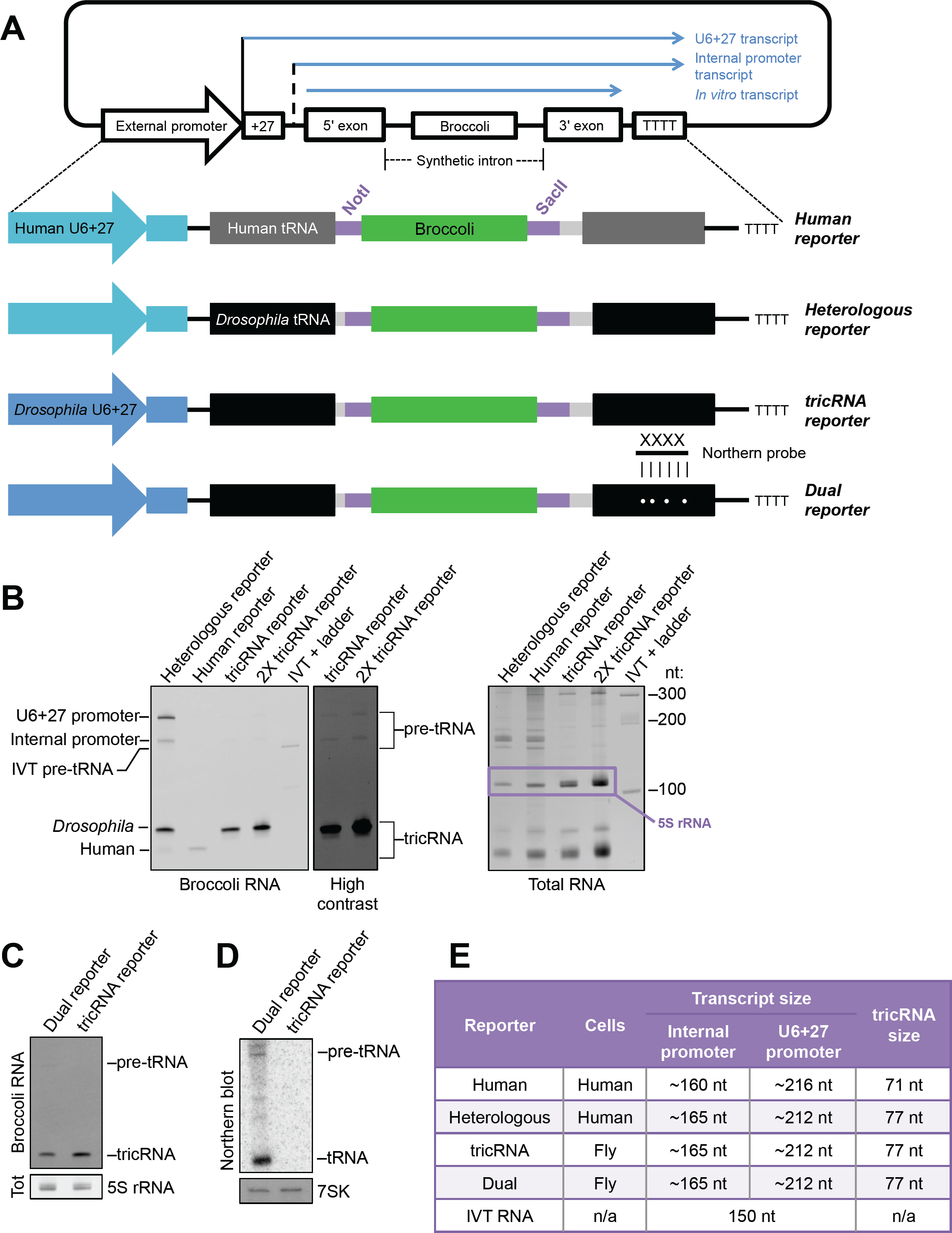
Developing a series of reporters. (A) Schematic of reporter constructs used in this study. Each contains either the human (lighter blue) or *Drosophila* (darker blue) external U6 snRNA promoter as well as the first 27 nucleotides of U6 snRNA (denoted as U6+27). The reporters also contain either a human (dark gray) or *Drosophila* (black) tyrosine tRNA. Additionally, each reporter contains the Broccoli fluorescent aptamer (green) as part of a synthetic intron. For details on the sequence and placement of the synthetic intron, see Supplementary Figure S1. The “dual reporter” contains four point mutations in the 3′ exon that allow specific detection by a Northern blot probe. (B) Reporter expression can be monitored by DFHBI stain. Left: DFHBI-stained gel showing expression of the heterologous, human, and tricRNA reporters. The top bands are pre-tRNA species, while the bottom bands are the tricRNA. Right: The gel was next stained with EtBr to detect total RNA. The 5S rRNA band is highlighted. (C) DFHBI-stained gel of the dual and tricRNA reporters. The 5S band from the EtBr-stained gel is shown as a loading control. (D) Northern blot of the dual and tricRNA reporters. 7SK is shown as a loading control. (E) Table showing the cell species used for each reporter, as well as the sizes of transcripts and tricRNAs.

To test expression of the reporters, we transfected the constructs into the appropriate cell line (Figure 2E) and harvested RNA for analysis. We visualized tricBroc using an in-gel fluorescence assay, described above and in (8, 20). Expression of tricBroc from the human reporter construct was relatively low, but the heterologous and tricRNA reporters express tricBroc at similar levels (Figure 2B, left). We also observed that the pre-tRNA bands are more prominent with the heterologous reporter (Figure 2B, left). This result could be due to higher transfection efficiency in human cells, more efficient processing of the construct in fly cells, better folding of the heterologous reporter pre-tRNA, or some combination of the three. To demonstrate that the pre-tRNA bands are, in fact, detectable following tricRNA reporter expression in fly cells, we also loaded twice the amount of total RNA in the adjacent lane. The pre-tRNA bands were only visible from the tricRNA reporter upon increasing the contrast of the image (Figure 2B, “High contrast”). Following Broccoli imaging, we re-stained the gel with EtBr to detect total RNA (Figure 2B, right). In all subsequent experiments, we cropped the EtBr-stained gel to show the 5S rRNA band as a loading control.

In order to detect both newly synthesized tricRNAs and tRNAs, we introduced four point mutations within the 3′ exon of the tRNA that allow specific detection of mature reporter tRNAs by northern blotting (Figure 2A, “dual reporter”). To test expression of these constructs, we transfected the dual and the tricRNA reporters into S2 cells. We visualized tricRNA expression using the in-gel staining assay described above (Figure 2C). To demonstrate that the dual reporter can be used to detect only the newly synthesized tRNAs, we performed northern blotting with a radiolabeled oligoprobe specific for the dual reporter tRNA. See Supplementary Figure S1 for details on the tRNA and the probe. As shown in (Figure 2D), the probe uniquely binds to the dual reporter tRNAs, and does not cross-hybridize to the tricRNA reporter transcript or to the endogenous tRNAs.

The reporter constructs described above each contain a stretch of 11 base pairs situated between the tRNA domain and the Broccoli domain (Figure 3A, left). To determine if there was a minimum length requirement for this fully-paired stem, we systematically deleted base pairs within the heterologous reporter, and transfected the resulting constructs into HEK293T cells. (Figure 3A, right). We found that reducing the stem below 7 bp in length inhibited tricRNA formation, whereas stems of 8 bp or greater yielded similar levels of tricBroc. Thus, we conclude that a stem of at least 8 bp is necessary for proper reporter function.

**Figure 3:**
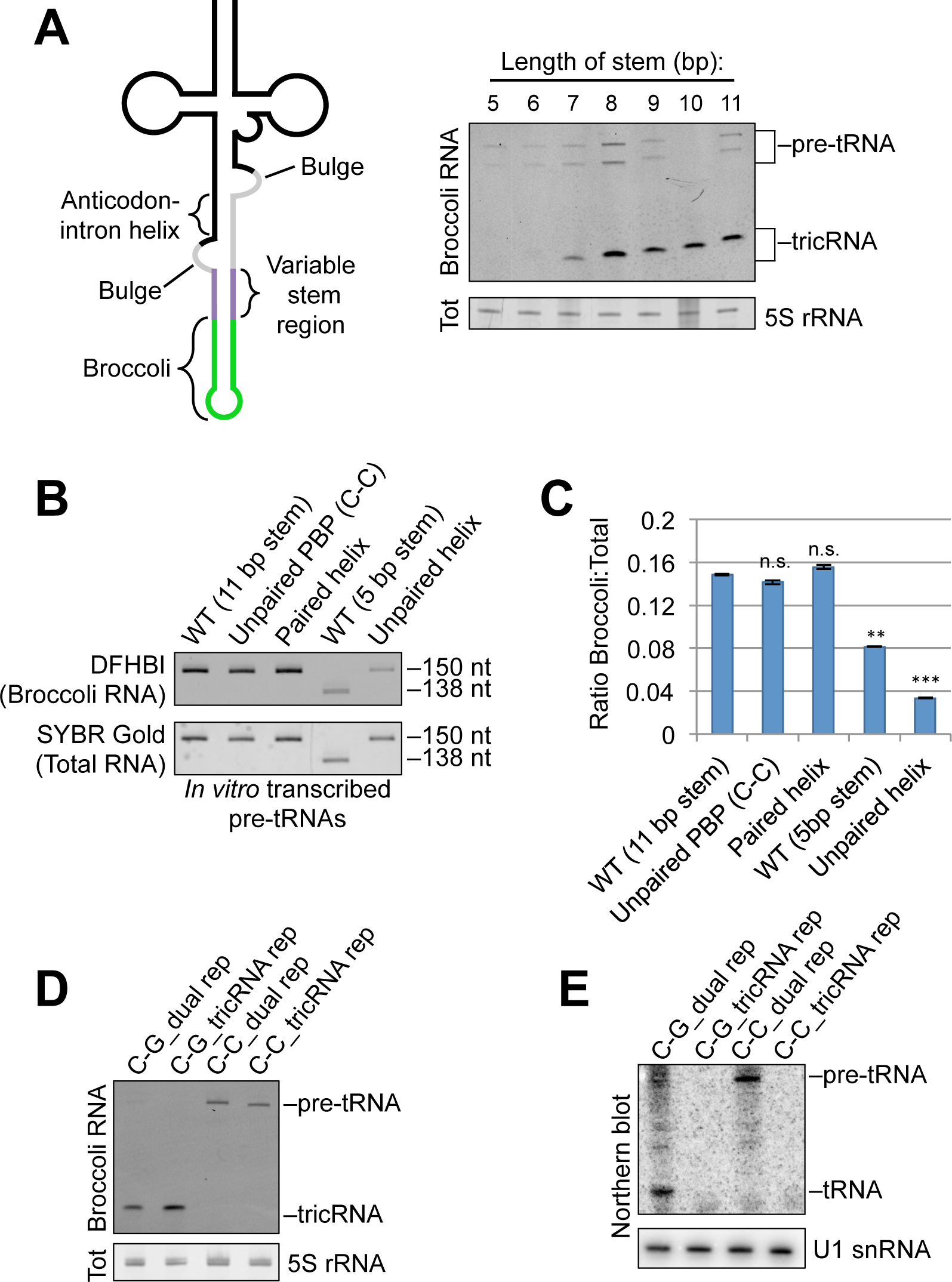
Characterization of reporters. (A) A stem of at least 8 base pairs is necessary for proper splicing of the heterologous tricRNA reporter. Left: Schematic of the reporter pre-tRNA, showing a variable stem length region. Right: In-gel fluorescence assay showing optimal stem length for the reporter. The 5S rRNA band from the total RNA gel was used as a loading control. (B) Mutations in the pre-tRNA can affect its folding. Top: DFHBI-stained gel of *in vitro* transcribed pre-tRNAs from various mutant constructs. Bottom: The gel was then stained in SYBR Gold to detect total RNA. (C) Quantification of technical duplicates from panel (B). Error bars denote standard error of the mean. **p<0.01, ***p<0.001, student’s t-test. (D) DFHBI-stained gel of the dual and tricRNA reporters, with and without an unpaired PBP (see Figure 4 for details). The 5S band from the EtBr-stained gel is shown as a loading control. (E) Northern blot of the dual and tricRNA reporters with and without an unpaired PBP. U1 snRNA is shown as a loading control.

To determine if the relative lack of tricBroc signal from the 5 bp stem construct was due to impaired folding of its pre-tRNA (and thus less efficient splicing), we generated *in vitro* transcripts from both the WT 5 bp and 11 bp ‘variable’ stem constructs and measured their fluorescence in replicate gels using the Broccoli staining assay (Figure 3B, compare lane 1 to lane 4). Note, an analysis of the three other *in vitro* transcripts in Figure 3B is described below as part of Figure 4. We also stained the gels with SYBR Gold to detect total RNA levels. Upon quantification, we found that compared to the 11 bp stem, the pre-tRNA from the 5 bp stem construct displayed significantly reduced fluorescence (Figure 3C), indicating that it may adopt folding conformations that can affect downstream processing. Therefore, all subsequent processing experiments utilized reporters with an 11 bp stem.

**Figure 4:**
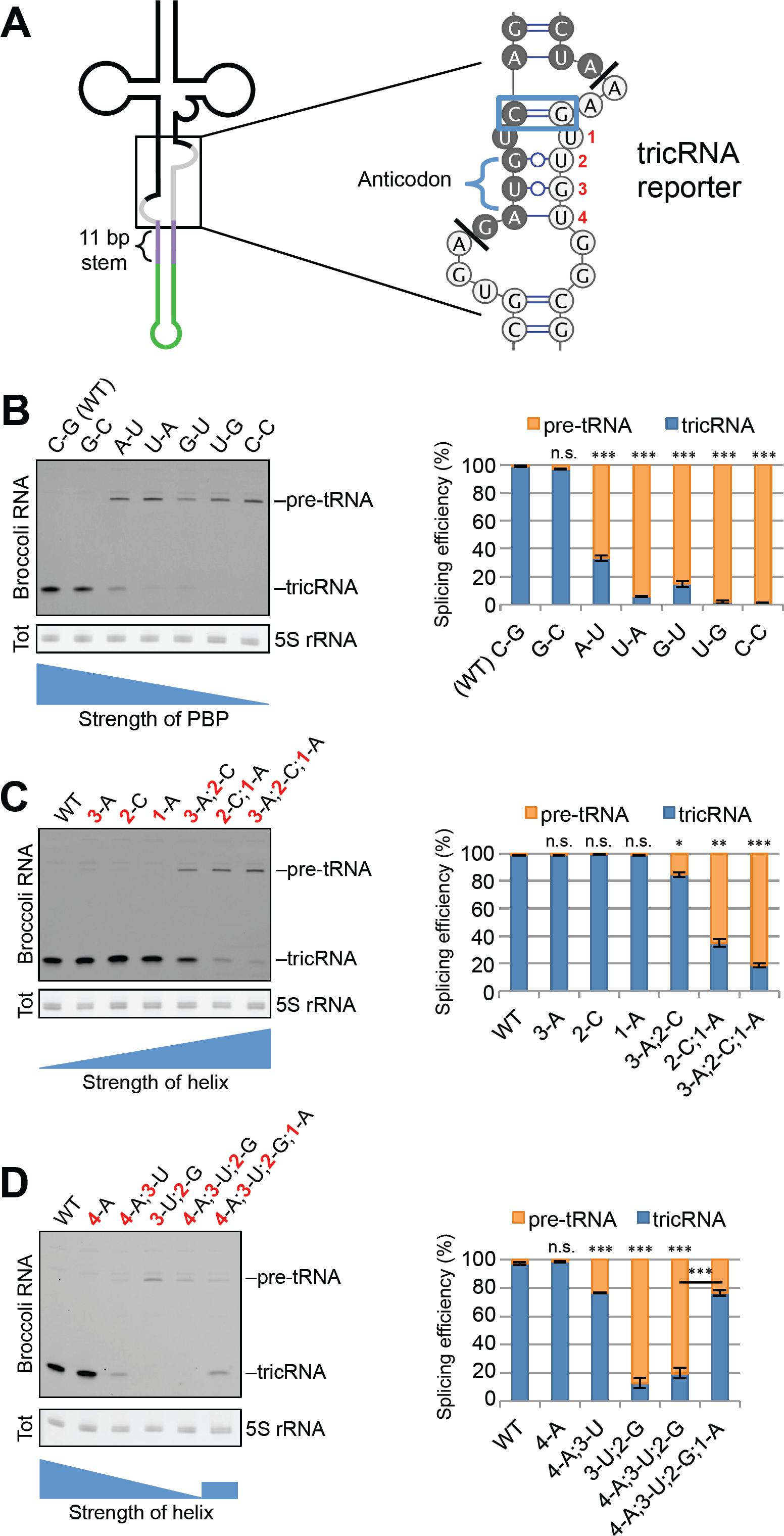
The helix portion of the BHB-like motif is important for proper tricRNA splicing. (A) Left: Schematic of the tricRNA reporter pre-tRNA, with emphasis of the BHB-like motif. Right: Nucleotide sequence of the BHB-like motif in the tricRNA reporter. The proximal base pair (PBP) is outlined in blue. The bases on the 3′ side of the helix are numbered 1-4. The darker bases are part of the tRNA, and the lighter bases are part of the intron. (B) Alterations to the PBP disrupt tricRNA splicing. Left: In-gel fluorescence assay of mutations to the PBP. Each lane represents a different identity of the PBP. The 5S rRNA band from the EtBr-stained gel is shown as a loading control. Right: quantification of three biological replicates. The pre-tRNA and tricRNA bands were expressed as a fraction of the total. Error bars denote standard error of the mean. ***p<0.001, student’s t-test. (C) Pairing the helix reduces tricRNA formation. Left: In-gel fluorescence assay of constructs pairing the helix. Each lane represents base pairs being introduced at various positions in the helix (see number positioning in panel A). The 5S rRNA band from the EtBr-stained gel is shown as a loading control. Right: quantification of three biological replicates. Quantifications were carried out as above. Error bars denote standard error of the mean. *p<0.05, **p<0.01, ***p<0.001, student’s t-test. (D) Unpairing the helix inhibits tricRNA production. Left: In-gel fluorescence assay of constructs unparing the helix. Each lane represents bases being paired or unpaired at various positions in the helix (see number positioning in panel A). The 5S rRNA band from the EtBr-stained gel is shown as a loading control. Right: quantification of three biological replicates. Quantifications were carried out as above. Error bars denote standard error of the mean. ***p<0.001, student’s t-test.

### The helix portion of the BHB-like motif is important for proper processing

We next focused on the structural features of pre-tRNAs. In archaea, the pre-tRNA contains a structural motif known as a bulge-helix-bulge (BHB), which consists of a 4 bp duplex flanked on each 3′ end by a single-stranded 3 nt bulge. This motif is both necessary and sufficient for splicing in archaea (21, 22). In contrast, eukaryotic splicing enzymes recognize the tRNA structure and although a motif resembling a BHB is often present, the structural requirements for splicing do not seem to be as rigid (23, 24). Therefore, we refer to this structure in our reporters as the “BHB-like motif” (Figure 4A). Due to its high degree of conservation among intron-containing tRNAs, we were interested in the first base pair of the anticodon-intron helix. Previous studies have referred to this feature as the A-I base pair (25); to avoid confusing this abbreviation with the bases Adenosine and Inonsine, we will refer to it as the proximal base pair (PBP). In all 16 of the intronic tRNAs in *Drosophila*, this base pair is a C-G (Supplementary Figure S6). To determine the importance of the PBP for proper pre-tRNA splicing and tricRNA formation, we generated a series of mutations in the tricRNA reporter and transfected the constructs into *Drosophila* S2 cells. To measure splicing efficiency, we performed an in-gel fluorescence assay on total RNA from the transfections (Figure 4B, left). We then expressed the intensities of the pre-tRNA and tricRNA bands as a fraction of the total (Figure 4B, right). Interestingly, we observed that switching the base pair to a G-C had little effect; however, weakening it to an A-U or U-A reduced tricRNA levels. Further weakening the helix to a G-U or U-G yielded almost no product, whereas unpairing it with a C-C mismatch resulted in complete inhibition of splicing. Notably, the fluorescence properties of the unpaired PBP (C-C) mutant pre-tRNA are equivalent to those of the WT (Figure 3B, compare lanes 1 and 2, quantification in Figure 3C); thus we conclude that folding of the aptamer tag is not perturbed by mutation of the PBP.

To determine if mutations that disrupt production of the tricRNA also affect expression of the mature tRNA, we mutated the PBP in the dual reporter construct, allowing analysis of both newly synthesized tRNAs and tricRNAs. As expected, unpairing the PBP affected production of both the tRNA and the tricRNA (Figures 3D and 3E). On the basis of these results, we conclude that the strength, rather than the identity, of the PBP is important for efficient biogenesis of the reporter tRNA and tricRNA.

The pre-tRNA encoded by the tRNA:Tyr reporter construct is predicted to contain several weak (or perhaps non Watson-Crick) base pairs within the anticodon-intron helix (Figure 4A, right). To determine if these weaker pairs are important, we first strengthened the helix by mutating the intronic bases (i.e. those on the 3′ side of the stem). Introducing a single base pair at any position within this helix had little effect on splicing efficiency (Figure 4C, compare lane 1 to lanes 2-4). However, introducing two or three base pairs at a time reduced tricRNA production (lanes 5-7). Importantly, the *in vitro* transcribed pre-tRNA containing a fully-paired helix displayed comparable fluorescence levels to that of the wild-type (Figure 3B, compare lanes 1 and 3, quantified in Figure 3C). The 5′ side of the helix is located in exon 1 of the tRNA and the 3′ side is part of the tricRNA; these halves must come apart in order to carry out exon ligation and intron circularization. Thus, we infer that if the helix is too tightly paired, splicing efficiency is reduced.

Mutations that are predicted to disrupt pairing within the anticodon-intron helix also inhibited formation of tricRNAs. We found that the integrity of the PBP alone is not sufficient for proper splicing; unpairing all of the other bases in the helix reduced tricRNA formation (Figure 4D, compare lanes 1 and 5). One possible explanation for this result is that this particular mutant pre-tRNA exhibits significantly reduced fluorescence compared to the wild-type (Figure 3B, compare lanes 1 and 5, quantified in Figure 3C), indicating that it might not fold properly and thus is inefficiently spliced. Similarly, fully unpairing the weak G•U pairs at positions 2 and 3 in the middle of the helix inhibited tricRNA production as well (lanes 3 and 4). However, introducing a single base pair adjacent to the proximal one in an otherwise unpaired helix partially rescued splicing (Figure 4D, lane 6). Interestingly, unpairing the distal base pair of the helix had no effect tricRNA splicing (Figure 4D, compare lanes 1 and 2). Because this helix base pair is furthest away from the proximal one, perhaps it may be less important for defining a specific structural motif. We conclude that a baseline level of helix pairing is required for optimal reporter tricRNA processing (see Discussion).

To elucidate whether these *cis*-element splicing determinants are applicable to other intron-containing pre-tRNAs, we developed a second reporter derived from *CR31143*, a *Drosophila* intron-containing tRNA:Leu_CAA_ gene (Supplementary Figure S2). Similar to our previous work with the tyrosine tricRNA reporter (9), we were able to increase tricRNA expression by using the *Drosophila* U6+4 (dU6) and U6+27 (dU6+27) external RNA polymerase III promoters (Supplementary Figure S2B). Although the BHB-like motif of the leucine reporter is distinct from that of the tyrosine reporter (Supplementary Figure S2A), we observed the same trends. Unpairing the PBP inhibited tricRNA production, and extending the helix by introducing two additional A-U base pairs reduced tricRNA splicing (Supplementary Figure S2C). Interestingly, we noticed that tricBroc RNA from the leucine reporter runs as a doublet rather than as a single band (Supplementary Figures S2B, S2C). We also found that extending the helix reproducibly affected migration of the doublet (Supplementary Figure S2C). To determine if the mutations affected splice site choice, we generated cDNA from these experiments and sequenced the tricRNA junctions (for details on this method, see ref. (9). We were surprised to find that tricRNAs from both wild-type and mutant leucine reporters each contained the same splice junction sequences, despite the fact that they resolve as doublets and that the mutant RNAs migrate more slowly (Supplementary Figure S2D). This result suggests that the doublets are structural isomers, or perhaps contain differentially modified bases.

We were also curious to determine whether the *cis*-acting elements we identified in flies were also important for tricRNA production in human cells. Accordingly, we generated a series of mutations in the heterologous reporter (Figure 2A) and expressed these constructs in HEK293T cells. We found similar trends in human cells (Supplementary Figure S3) as we did in *Drosophila* (Figure 4). Unpairing the PBP (C-C) in the heterologous reporter inhibited tricRNA production and resulted in a buildup of pre-tRNA precursors. Furthermore, introducing additional base pairs within the helix was inversely proportional to reporter tricRNA formation (Supplementary Figure S3). Taken together, these data reveal that similar rules govern tricRNA biogenesis in both vertebrates and invertebrates.

### Pre-tRNA cleavage in *Drosophila* is carried out by orthologs of the TSEN complex

After characterizing *cis* elements important for proper tricRNA splicing, we next sought to identify tricRNA processing factors. We searched for *Drosophila* homologs of known human tRNA processing factors and found candidates for many of these genes (Figures 5A, 6A). To analyze these factors, we developed an RNA interference (RNAi) assay in S2 cells. Candidate processing factors were depleted using dsRNA for seven days, at which point we transfected the dual reporter into these cells. We continued applying dsRNA daily to continue the knockdown, and cells were harvested three days post transfection for RNA analysis.

**Figure 5:**
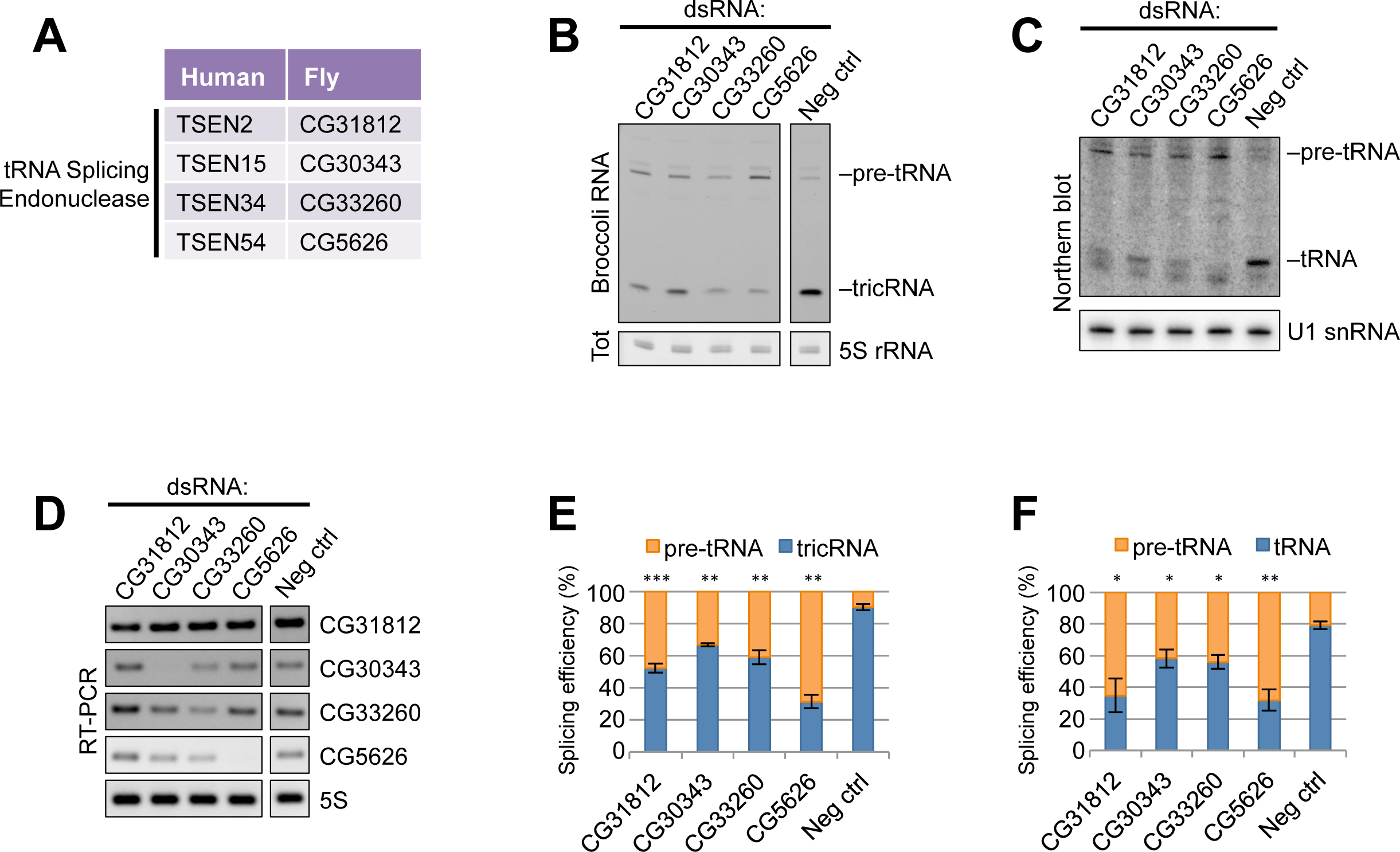
Knocking down any member of the TSEN complex inhibits reporter tricRNA and tRNA production. (A) Table of human tRNA processing factors and *Drosophila* sequence homologs. (B) In-gel fluorescence assay of RNA from S2 cells depleted of TSEN complex members. The pre-tRNA and tricRNA bands are identified. The 5S rRNA band from the EtBr-stained gel is shown as a loading control. (C) Northern blot of RNA from S2 cells depleted of TSEN complex members using a probe specific to the dual reporter. The pre-tRNA and tRNA bands are identified. U1 snRNA is used as a loading control. (D) RT-PCR for TSEN complex members to test knockdown efficiency in S2 cells. RT-PCR for 5S rRNA was used as a control. (E) Quantification of three biological replicates of (B). Error bars denote standard error of the mean. ***p<0.001, **p<0.01, student’s t-test. (F) Quantification of three biological replicates of (C). Error bars denote standard error of the mean. **p<0.01, *p<0.05, student’s t-test.

We first investigated potential members of the tRNA splicing endonuclease complex (Figure 5A), which cleaves pre-tRNAs in both fungi (where it is called SEN) and humans (called TSEN) (26). Depletion of any member of this complex (Figure 5D) results in a dramatic reduction in both reporter tRNA and tricRNA formation (Figures 5B and 5C). Concordantly, knockdown of these proteins also results in accumulation of the pre-tRNA, visible as the top band in the Broccoli stained gel (Figure 5B). The pre-tRNA band can also be detected by northern blotting, as the probe binds to a region in the 3′-exon (Figure 5C). The tricRNA and tRNA results are quantified in Figures 5E and 5F, respectively. From these data, we conclude that these genes indeed perform the functions of the TSEN complex in *Drosophila*.

### *Drosophila* tRNA intron circularization proceeds through the direct ligation pathway

In archaea and eukarya, cleavage of an intron-containing pre-tRNA yields non-canonical RNA ends: a 5′-OH and a 2′,3’-cyclic phosphate. Although plants and fungi use a roundabout “healing and sealing” approach, archaeal and animal cells are thought to utilize the direct ligation pathway, wherein there is no need for addition of an external phosphate (27). Previously, we showed that knockdown of CG9987 (now called RtcB) in *Drosophila* larvae and pupae resulted in a significant decrease in *tric31905* levels, along with a decrease in the corresponding tRNA:Tyr (8). However, this approach could not distinguish between newly-transcribed and pre-existing RNAs, and therefore likely underestimated the effects of RtcB depletion. Accordingly, we used the S2 cell RNAi system to assess levels of newly synthesized tRNAs and tricRNAs during RtcB knockdown. We also analyzed cells depleted of the other members of the human tRNA ligase complex, Archease and Ddx1 (Figures 6A and 6D). Depletion of RtcB caused a significant decrease in the levels of both the mature reporter tRNA as well as its corresponding tricRNA (Figures 6B and 6C; quantified in Figures 6E and 6F). Similarly, depletion of Archease and Ddx1 also resulted in reduction of both reporter RNAs. Importantly, depletion of the tRNA ligase complex did not cause accumulation of the pre-tRNA, reaffirming previous findings that cleavage and ligation are separable processes (28, 29). We conclude that RtcB is the *Drosophila* tRNA and tricRNA ligase, and that its activity is regulated *in vivo* by Archease and Ddx1.

**Figure 6:**
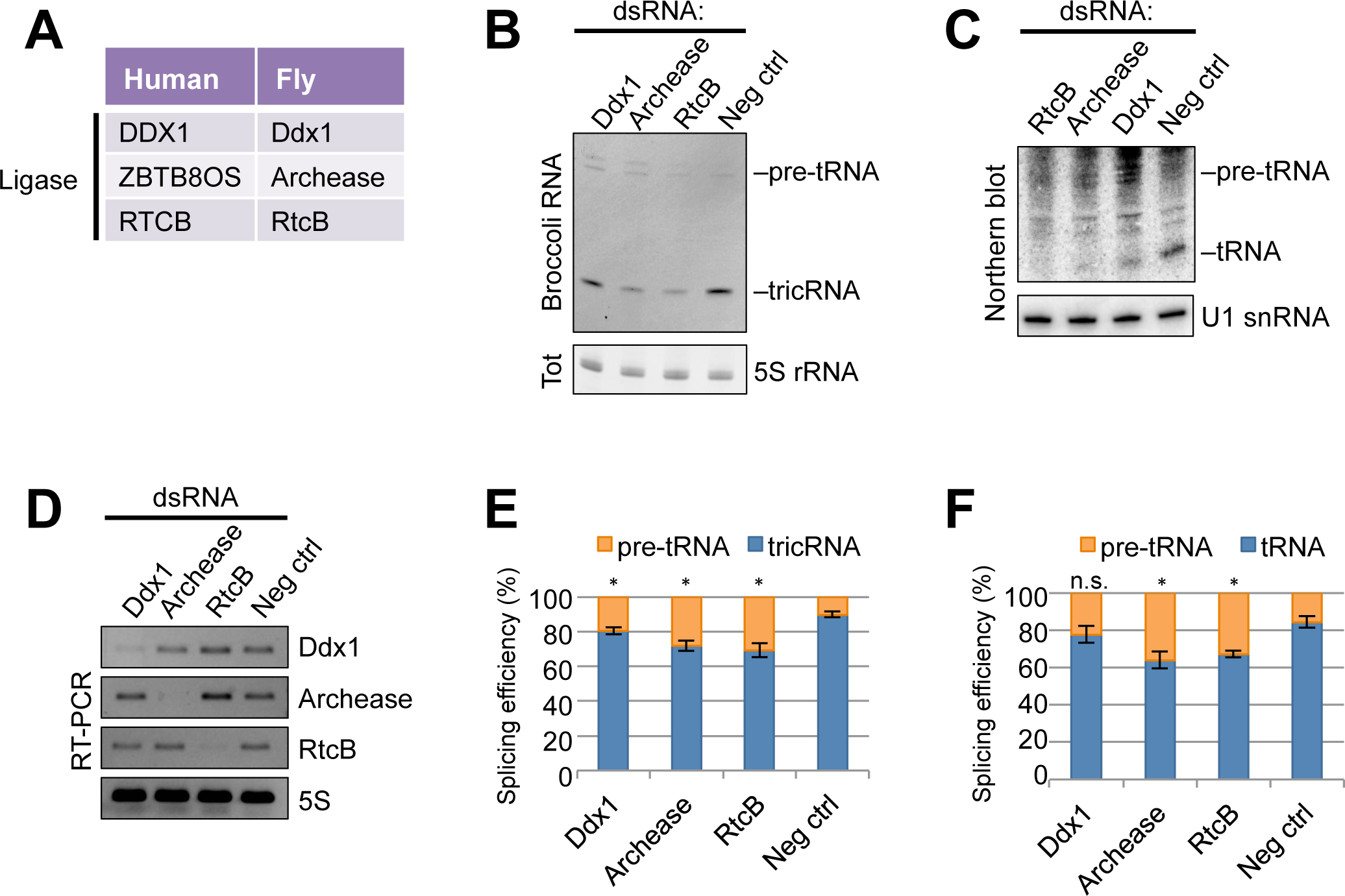
Knocking down RtcB ligase or its associated factors Archease and Ddx1 inhibits reporter tricRNA and tRNA production. (A) Table of human tRNA processing factors and *Drosophila* sequence homologs. (B) In-gel fluorescence assay of RNA from S2 cells depleted of RtcB, Archease, or Ddx1. The pre-tRNA and tricRNA bands are identified. The 5S rRNA band from the EtBr-stained gel is shown as a loading control. (C) Northern blot of RNA from S2 cells depleted of RtcB, Archease, or Ddx1 using a probe specific to the dual reporter. The pre-tRNA and tRNA bands are identified. U1 snRNA is used as a loading control. (D) RT-PCR for RtcB complex members to test knockdown efficiency in S2 cells. RT-PCR for 5S rRNA was used as a control. (E) Quantification of three biological replicates of (B). Error bars denote standard error of the mean. *p<0.05, student’s t-test. (F) Quantification of three biological replicates of (C). Error bars denote standard error of the mean. *p<0.05, student’s t-test.

### A *Drosophila* ortholog of TRPT1 does not participate in tricRNA biogenesis

While searching for tRNA processing factors, we found that the human genome contains a homolog of yeast Tpt1, the phosphotransferase that participates in the healing and sealing pathway (See discussion for more details). To determine if the *Drosophila* ortholog, CG33057 (Supplementary Figure S4A), participates in tricRNA biogenesis, we used the S2 cell RNAi assay described above. Depletion of CG33057 (Supplementary Figure S4D) resulted in no measurable difference in tricRNA levels (Supplementary Figure S4B, quantified in Supplementary Figure S4C), indicating that this protein is unlikely to play a role in *Drosophila* tRNA intron circularization.

### tricRNA turnover is initiated by an endoribonuclease

Due to their lack of free 5′ and 3′ ends, circular RNAs exhibit a stability not afforded to their linear counterparts (8, 30, 31). Passive degradation of broken circles likely occurs via exonucleolytic decay, however active degradation of tricRNAs must be initiated by some type of endonuclease. To determine which enzymes are responsible for tricRNA turnover, we depleted S2 cells of candidate endoribonucleases, excluding those known to be directly involved in the RNAi pathway. We chose Zucchini, which is involved in piRNA biogenesis (32); Smg6, which is part of the nonsense-mediated mRNA decay pathway (33, 34); Dis3, a dual endo/exonuclease component of the nuclear exosome (35, 36); and Clipper, which has been shown to cleave RNA hairpins and is a homolog of mammalian CPSF 30K subunit (37, 38). Following knockdown (Figure 7C) and subsequent RNA isolation, we assessed endogenous tricRNA abundance by northern blotting for *tric31905*. As shown in Figure 7A and quantified in Figure 7B, depletion of Zucchini, Smg6, or Dis3 had little to no effect on *tric31905* levels, whereas depletion of Clipper resulted in a modest increase in *tric31905*. We also performed a double knockdown of Dis3 and Clipper and observed a similar increase in *tric31905* levels (Figures 7A and 7B). In contrast, overexpression of FLAG-tagged Dis3 or Clipper did not reduce levels of *tric31905* (Figure 7D). Taken together, our data suggest that turnover of tric31905 is in part initiated by Clipper activity.

**Figure 7:**
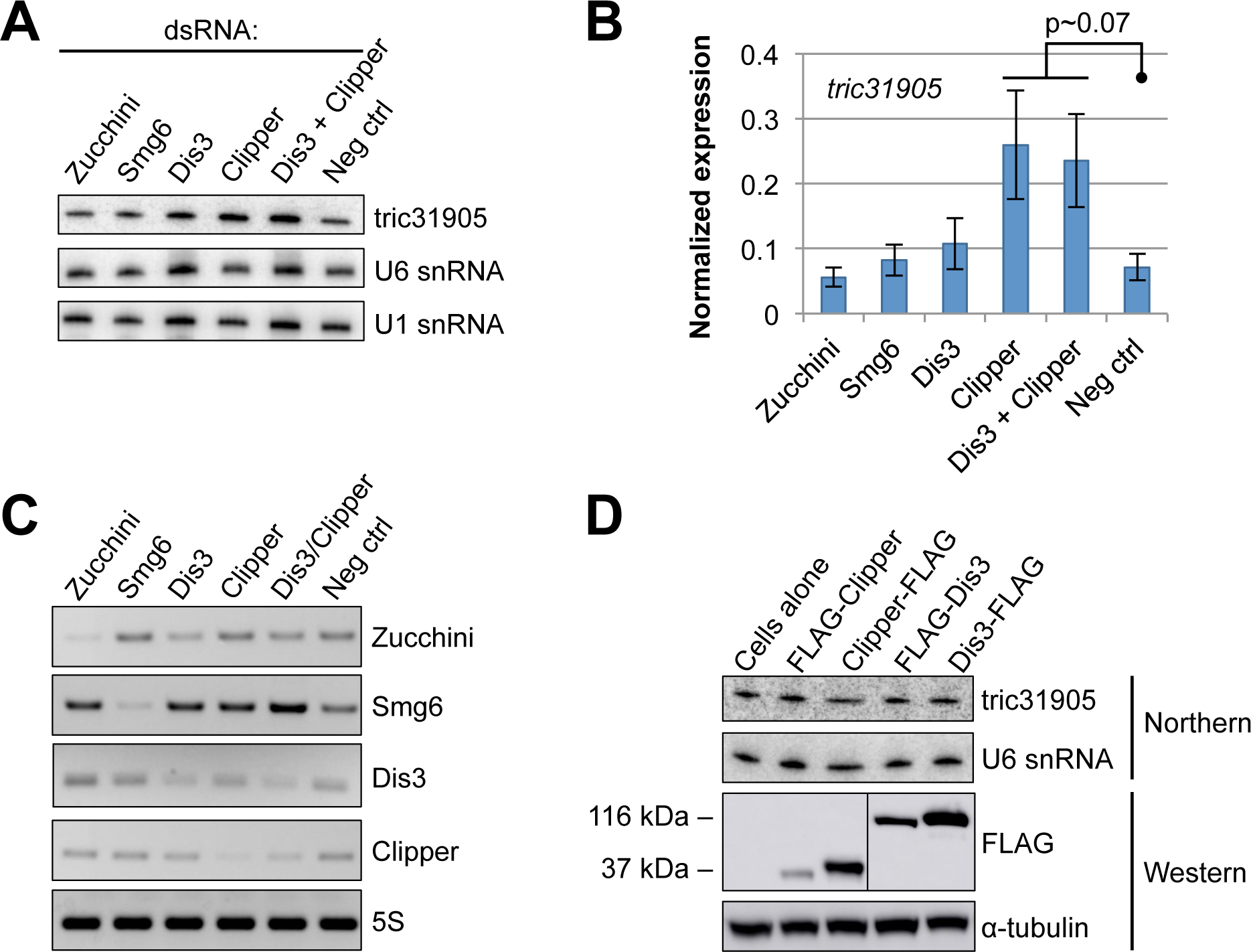
Clipper endonuclease is required for tricRNA turnover. (A) Northern blot of RNA from endonuclease-depleted S2 cells. U1 and U6 snRNA are shown as loading controls. (B) Quantification of three biological replicates. The tric31905 bands were normalized to U6 snRNA bands and average values were plotted. Error bars denote standard error of the mean. The p-value was calculated using student’s t-test. (C) RT-PCR for candidate endonucleases to test knockdown efficiency in S2 cells. RT-PCR for 5S rRNA was used as a control. (D) Western blot showing overexpression of candidate endonucleases. S2 cells were transfected with FLAG-tagged endonuclease constructs. Cells from each condition were split in half for both RNA and protein isolation. The top two panels are a Northern blot of RNA from the experiment, and the bottom two panels are a Western blot of protein lysate from the experiment. For the Northern, U6 snRNA is used as a loading control; for the Western, tubulin is used as a loading control.

### RtcB replacement yeast generate tricRNAs

A unique aspect of eukaryotic pre-tRNA processing is the differential fate of the intron in various organisms. The genomes of plants and fungi do not contain RtcB homologs and their tRNA introns are typically linear (6, 39). In contrast, archaeal and metazoan genomes contain RtcB and they express tricRNAs (7, 8). Interestingly, bacterial *RtcB* can complement deletion of yeast *Rlg1/Trl1*, which is an essential gene (11). To determine if “RtcB replacement yeast” can make tricRNAs, we isolated total RNA from a variety of control and mutant yeast strains and performed RT-PCR to analyze expression of an intronic tRNA:Ile_UAU_ gene. Following ‘rolling circle’ reverse transcription, moderately sized circular RNA templates like tricRNAs are known to produce linear cDNAs that contain many tandem copies of the sequence (Figure 8A). This concatamer has multiple binding sites for primers, thus producing a ladder of PCR products with predictable sizes (9). Linear RNA templates do not produce such ladders. As shown, RT-PCR products from yeast expressing *RtcB* display a ladder of bands with the predicted periodicity (Figure 8B, lanes 2, 4 and 5). Thus, RtcB is active even in the presence of Rlg1/Trl1 (Figure 8B, lane 2). Moreover, linear tRNA introns can serve as substrates for RtcB and this activity appears to be sufficient for tricRNA formation in yeast. As a control, we examined RNA from yeast lacking Xrn1 (Figure 8B, lanes 3 and 4), a 5′ to 3′ exonuclease that is known to degrade tRNA introns (6). In the *xrn1Δ* strain, linear tRNA introns accumulate (6) and can be detected by northern analysis.

**Figure 8:**
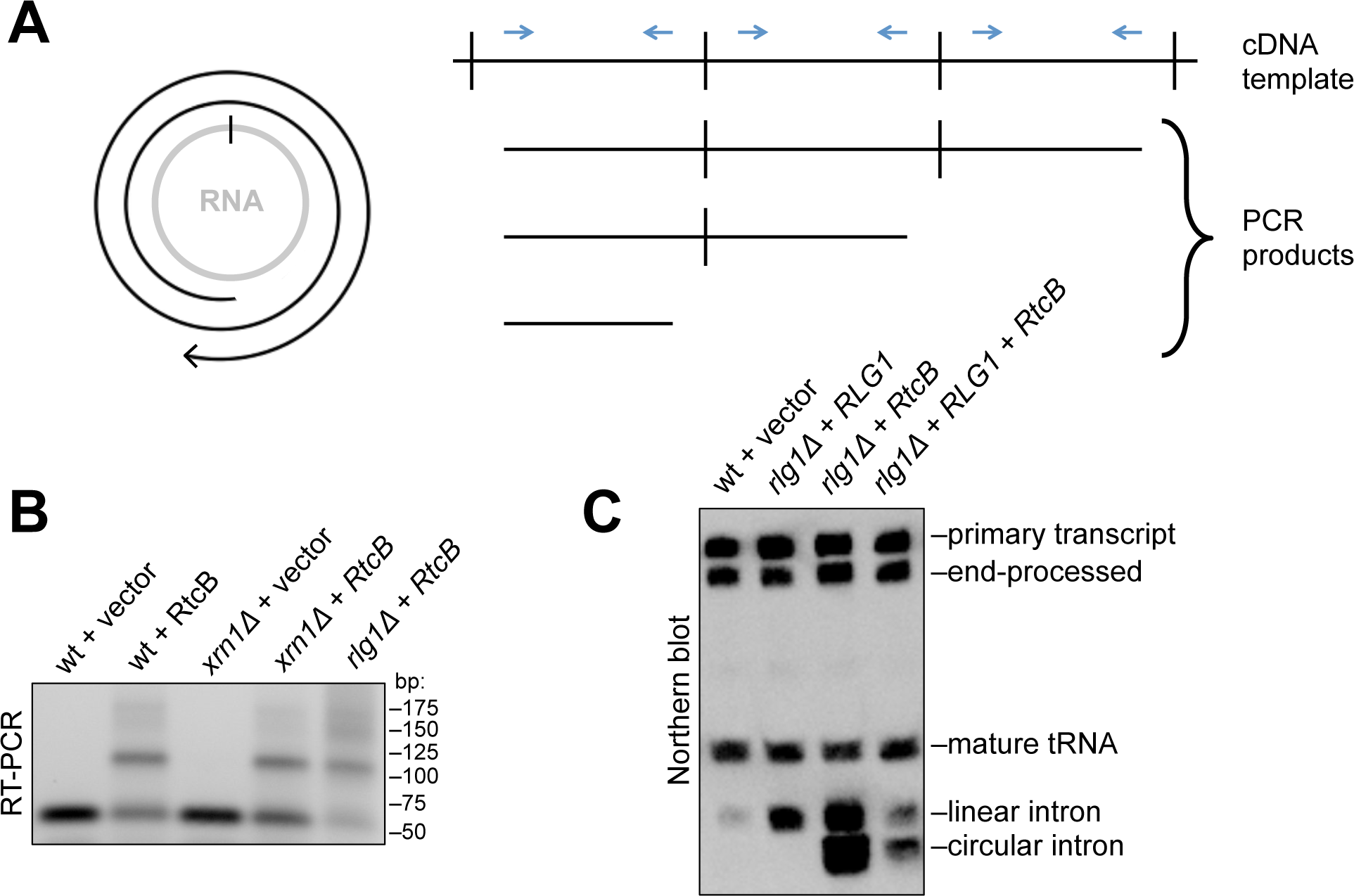
RtcB replacement yeast generate tricRNAs. (A) Left: schematic of cDNA generated from a circular template. Right: schematic of possible RT-PCR products. Reverse transcription of circRNAs can generate tandem copies of the circular sequence, particularly if the circle is relatively small. The resultant cDNA contains many binding sites for PCR primers. (B) RT-PCR for a tRNA intron yields a ladder of products only in yeast strains that express RtcB. (C) Northern blot of RNA from yeast bearing various deletions and enzyme replacements using a probe against both the tRNA:Ile_UAU_ 5′ exon and intron.

To confirm these findings, we performed northern blotting with probes targeting the intron and 5′ exon of tRNA:Ile_UAU_ (Figure 8C). In all cells expressing RtcB, we observed an anomalously migrating intron fragment consistent with the presence of a circle (Figure 8C, Supplementary Figure S5). Similar to the RT-PCR data above, this fragment is present in the RtcB replacement (rlg1Δ + RtcB) strain (Figure 8C) as well as in the absence of Xrn1 (Supplementary Figure S5A). To further confirm the circular nature of this new intron-specific fragment, we treated RNA from various yeast strains with terminator exonuclease (TEX) and performed a Northern blot using a probe against both the intron and 5′ exon of tRNA:Ile_UAU_. As shown in Supplementary Figure S5B, the circular intron is resistant to TEX cleavage. From these experiments, we conclude that RtcB replacement yeast produce circular introns.

## DISCUSSION

Removal of introns from pre-tRNAs is an essential step in tRNA maturation (26). Once thought to be discarded as waste products, we now know that metazoan tRNA introns are ligated into circular RNAs in animals such as *D. melanogaster* and *C. elegans*. Here, we demonstrate that specific *cis*-element structures are necessary for production of both tRNAs and tricRNAs. Moreover, we show that tricRNAs are produced by the tRNA splicing machinery in *Drosophila*, and tricRNA degradation is initiated by endonuclease activity. Finally, we provide evidence that the direct ligation pathway is the primary generator of tricRNAs in *Drosophila*, and that RtcB is the factor that determines tRNA intron circularization in eukaryotes.

### Structure and stability of pre-tRNAs

In archaea, intron-containing pre-tRNAs and pre-rRNAs are known to contain a highly conserved BHB motif (40). For many years, this motif was thought to be strictly required for proper splicing, although more recent work indicates that certain pre-tRNAs contain more relaxed BHB structures (41). We have found that *Drosophila* also display a wide variety of BHB-like motifs (Supplementary Figure S6). Perhaps the eukaryotic pre-tRNA structure is more relaxed in order to facilitate TSEN substrate interaction. Despite this diversity, we invariably identified a 3 bp bulge on the proximal side of the anticodon-intron helix (i.e. at the cut site between the intron and the 3′ exon) (Supplementary Figure S6). Similarly, there is a constant number of bases between the 5′ base of the PBP and the last base of the 5′ exon (6 nt, including the C in the PBP; see Supplementary Figure S6). These regularities, combined with the lack of a strict BHB, suggest that *Drosophila* intron-containing pre-tRNAs are spliced in a manner akin to the ‘molecular ruler’ mechanism outlined by others (23, 24). Due to an overall dearth of high-resolution data in eukaryotes, we can only speculate with regard to the precise structure of the substrate-bound form of the metazoan TSEN complex.

Although we can draw broad conclusions from the various helix pairing and unpairing experiments, we cannot know for sure if these mutations cause alternative pre-tRNA structures to form. Similarly, we do not know the extent to which these mutations affect pre-tRNA stability. For example, we show that certain mutations in the pre-tRNA impair aptamer fluorescence *in vitro* (Figure 3B) and thus may affect RNA folding and downstream processing *in vivo*. Unpairing all of the distal bases in the anticodon-intron helix appeared to reduce stability of the pre-tRNA *in vivo* (Figure 4D). In contrast, there is a clear accumulation of the pre-tRNA in many other mutant constructs (Figures 4B and 4C). Perhaps mutation of the PBP also disrupts the structure of the invariant 3 nt bulge and thereby impairs recognition of the intron-3′ exon cut site. Similarly, if the helix region is too tightly paired, the rigidity of the duplex could interfere with positioning of the cut sites or otherwise reduce interaction with the TSEN complex. In the future, high-resolution (e.g. NMR) structure-function analysis of wild-type and mutant BHB-like motifs could shed light on these issues.

### Fungi and metazoa: distinct intron fates

Two pathways for tRNA ligation exist in nature: the “healing and sealing” pathway, utilized by plants and fungi, and the “direct ligation” pathway, found in archaea and animals (26). Plants and fungi lack RtcB-family enzymes and thus only have one way to ligate tRNA exons (42, 43). Similarly, archaea do not possess healing and sealing enzymes and therefore can only join RNA ends by direct ligation (27). Although metazoans appear to primarily use the direct ligation pathway (27), there is evidence for healing and sealing-type splicing in higher organisms. For example, mammalian genomes contain an RNA polynucleotide kinase (CLP1) and a 2′,3’-cyclic nucleotide phosphodiesterase (CNP) that can perform the corresponding activities of yeast Rlg1/Trl1 (44, 45); they also contain a putative 2′-phosphotransferase called TRPT1 (46). Additionally, an Rlg1/Trl1-like tRNA ligase activity has been detected in human cells (47). However, mouse *Cnp1* and *Trpt1* are not essential, as knockouts of either of these genes do not affect organismal viability (48, 49). Thus, a yeast-like tRNA splicing pathway may exist in mammals, but it likely contributes little to overall tRNA production. In this study, we have shown that direct ligation is the primary tRNA and tricRNA biogenesis pathway used in flies (Figures 5, 6, and Supplementary Figure S4). Similar to the situation in mammals, we cannot exclude a role for the *Drosophila* healing and sealing pathway in developmental regulation or in response to cellular stress. Perhaps this alternative pathway might be utilized in a tissue-specific or stress-induced manner. Additional studies will be needed to assess the overall contribution of this pathway to the regulation of tRNA and tricRNA expression in *Drosophila*.

It is interesting to consider why *Drosophila* tRNA introns are circularized, rather than being immediately degraded, as is their typical fate in yeast (6). One possibility is that tricRNAs serve an important function, and so production of these circRNAs would outweigh the energetic benefits of creating a supply of free nucleotides. Previously, we observed a high degree of conservation of *tric31905* among the twelve fully-sequenced Drosophilid species (8). In particular, there is significant co-variation within the predicted *tric31905* secondary structure, suggesting that there has been selection for a specific structure over evolutionary time (8). Consistent with these observations, we previously identified reads from small RNA-seq datasets that map to the intron of *CR31905* (8). Taken together with Figure 7, it seems likely that Clipper is involved in the observed downstream processing of *tric31905* into 21nt fragments (8). Whether these downstream processing events lead to formation of RNA silencing complexes has yet to be determined. Alternatively, endonucleolytic cleavage by Clipper might initiate the process of tricRNA degradation. Given that Clipper exhibits a preference for binding RNAs with G-and/or C-rich regions (38), it is also possible that different endonucleases may cleave different tricRNAs.

Irrespective of the ultimate fate or function of metazoan tRNA introns, these observations raise an important evolutionary question. As mentioned above, the fate of tRNA introns varies between different organisms, and the presence of tricRNAs appears to correlate with the occurrence of RtcB in the corresponding genome. Is RtcB the determining factor for tRNA intron circularization? Here, we showed that replacing a healing and sealing pathway enzyme in yeast with bacterial RtcB results in tricRNA formation. These data suggest that intron circularization may be a normal “byproduct” of RtcB exon ligation. RtcB is the eponymous member of an RNA ligase family that requires a 5′-OH and a 2′,3’-cyclic phosphate in order to carry out its enzymatic activity (29). Because these atypical RNA termini are generated by RNase L type endonucleases, a second prediction is that mRNA exons that are ligated by RtcB would also generate circular introns. Indeed, recent work strongly supports this idea. IRE1 is an RNase L family endonuclease that cleaves *HAC1* mRNA during the unfolded protein response in yeast (50) and *XBP1* mRNA in metazoan cells (51). The long-sought RNA ligase responsible for catalyzing this unconventional RNA splicing event during the metazoan UPR is RtcB (52–54). In budding yeast, Rlg1/Trl1 ligates the two *HAC1* exons and the linear intron is degraded. However, in the same *rlg1Δ+RtcB* yeast strain used here, the *HAC1* mRNA intron was also recently shown to be circularized (55). Altogether, these findings provide strong support for the notion that fungal tRNA introns are primarily linear because yeast lack RtcB.

## ACKNOWLEDGEMENT

The authors thank Dr. Steve Rogers for assistance with S2 cells, and Dr. Stewart Shuman for the RtcB plasmid and *rlg1Δ+RtcB* yeast strain.

## FUNDING

This work was supported by grants from the National Institutes of Health, R01-GM118636 (to A.G.M.), and R01-GM122884 (to A.K.H.). C.A.S. was supported in part by a National Science Foundation Graduate Research Fellowship, DGE-1650116, and by a Dissertation Completion fellowship from the University of North Carolina Graduate School. A.B. was supported in part by an Ohio State University Pelotonia Undergraduate Fellowship. Funding for open access charge: National Institute of General Medical Sciences.

## CONFLICT OF INTEREST

The authors declare no conflict of interest.

