## Supplementary Figures for "Molecular determinants of metazoan tricRNA biogenesis"

**A**

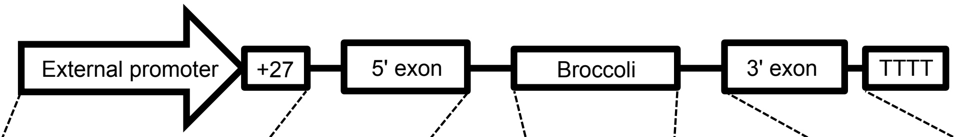


| | | | | | | | |
|---|---|---|---|---|---|---|---|
| **Human reporter** | pAV U6+27 (Addgene #25709) | TRYGTA3-1 | GCGGCCGC | Broccoli | GTGGCCGCGG | AGGT | TRYGTA3-1 |
| **Heterologous reporter** | pAV U6+27 (Addgene #25709) | CR31905 | AGTGCGGCCGC | Broccoli | GTGGCCGCGG | TGTTGAA | CR31905 |
| **tricRNA reporter** | snRNA:U6:96Ab | CR31905 | AGTGCGGCCGC | Broccoli | GTGGCCGCGG | TGTTGAA | CR31905 |
| **Dual reporter** | snRNA:U6:96Ab | CR31905 | AGTGCGGCCGC | Broccoli | GTGGCCGCGG | TGTTGAA | CR31905* |

ATCCATAGGTCGCCGGTTCGATTCCGGCTCGGAGGA

**B**

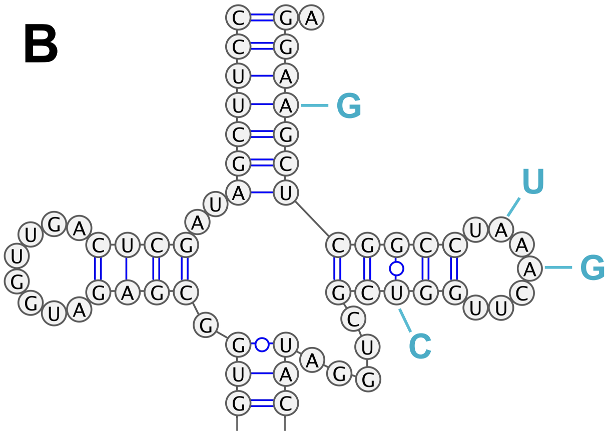


Supplementary Figure S1: Detailed description of reporters used in this study. (A) Table of the specific sequences used in each reporter. Restriction enzyme sites are in red. For the dual reporter, the entire 3' exon sequence is shown. The four mutations are shown in blue. The orange line shows where the probe binds. (B) Partial structure of the CR31905 tRNA molecule, indicating where the mutations were made.

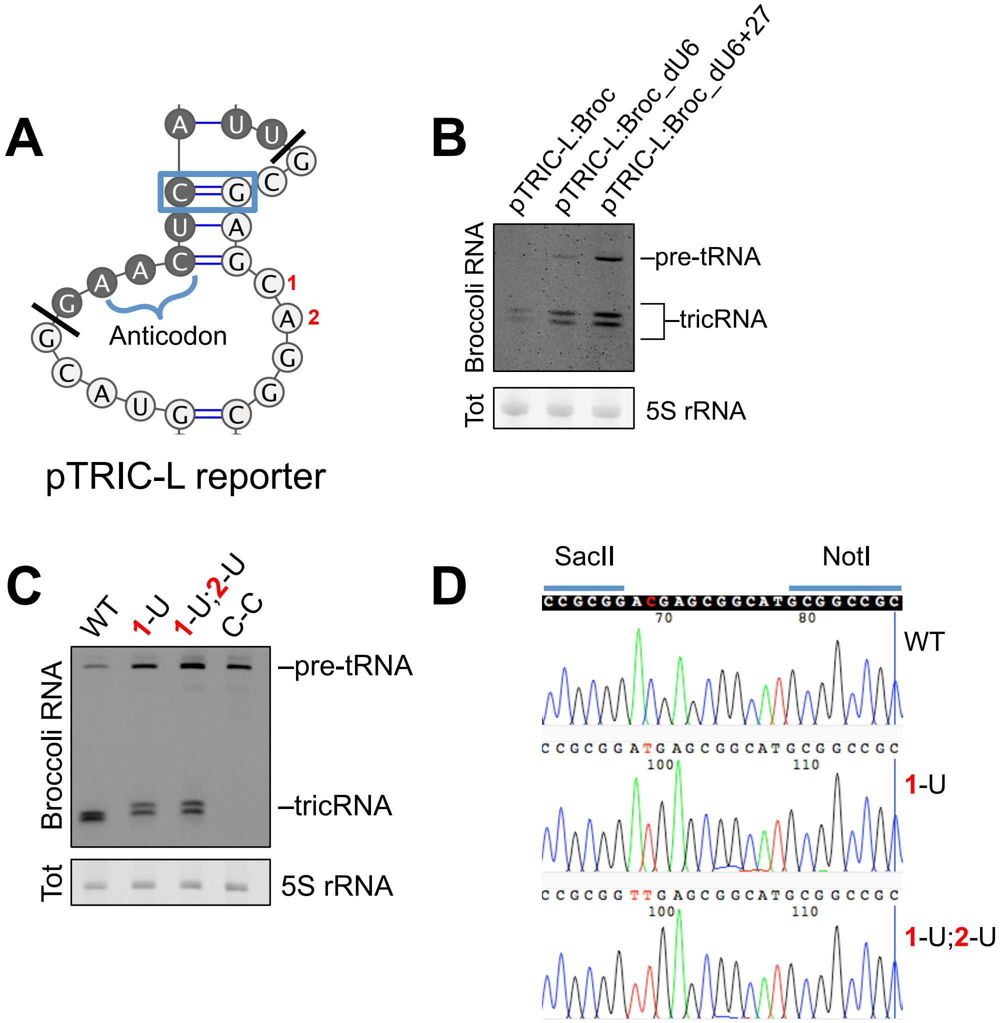


Supplementary Figure S2: Similar trends observed using a different tricRNA reporter. (A) Nucleotide sequence of the BHB-like motif in the pTRIC-L tricRNA reporter (see Fig. S5 for endogenous BHB-like motif of CR31143). The PBP is boxed in blue. Mutated residues are numbered 1 and 2. The darker shaded bases are part of the tRNA, and the lighter shaded bases are part of the intron. (B) In-gel fluorescence assay of RNA from S2 cells transfected with the pTRIC-L:Broc reporter to test expression of the U6 and U6+27 external RNA polymerase III promoters. The pre-tRNA and tricRNA bands are identified. The 5S rRNA band from the EtBr-stained gel is shown as a loading control. (C) In-gel fluorescence assay of RNA from S2 cells transfected with the wild-type and mutant reporters. The pre-tRNA and tricRNA bands are identified. The 5S rRNA band from the EtBr-stained gel is shown as a loading control. (D) Sequence traces of tricRNA junctions from the experiment in (C). The mutations introduced by site-directed mutagenesis (1-U and 1-U;2-U) can be seen in the traces.

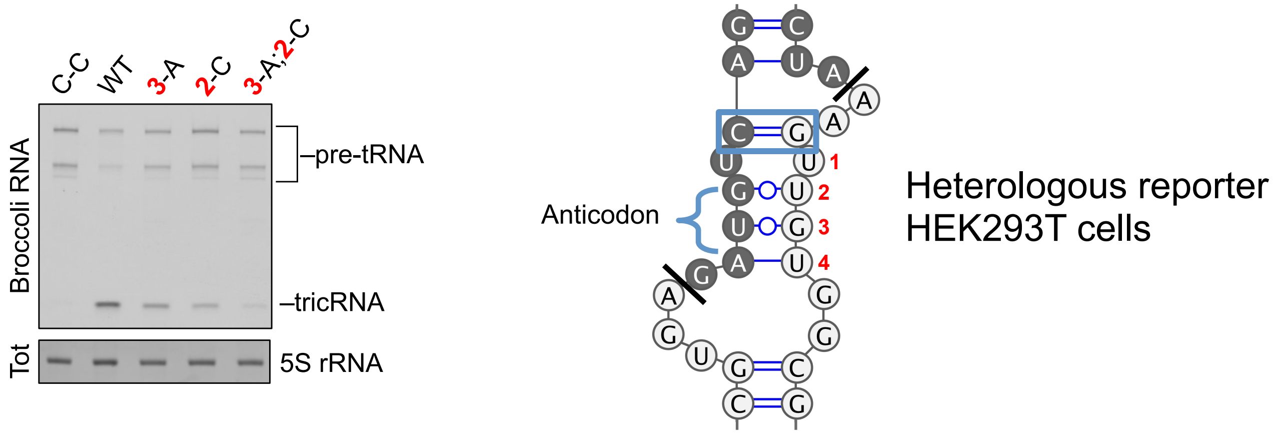


Supplementary Figure S3: mutations in the heterologous tricRNA reporter have similar effects on tricRNA splicing. Left: In-gel fluorescence assay of RNA from HEK293T cells transfected with the heterologous reporter to test expression of various mutant constructs. The pre-tRNA and tricRNA bands are identified. The 5S rRNA band from the EtBr-stained gel is shown as a loading control. Right: Nucleotide sequence of the BHB-like motif in the heterologous reporter. The PBP is outlined in blue. The other base pairs of the helix are numbered 1-4. The darker bases are part of the tRNA, and the lighter bases are part of the intron.

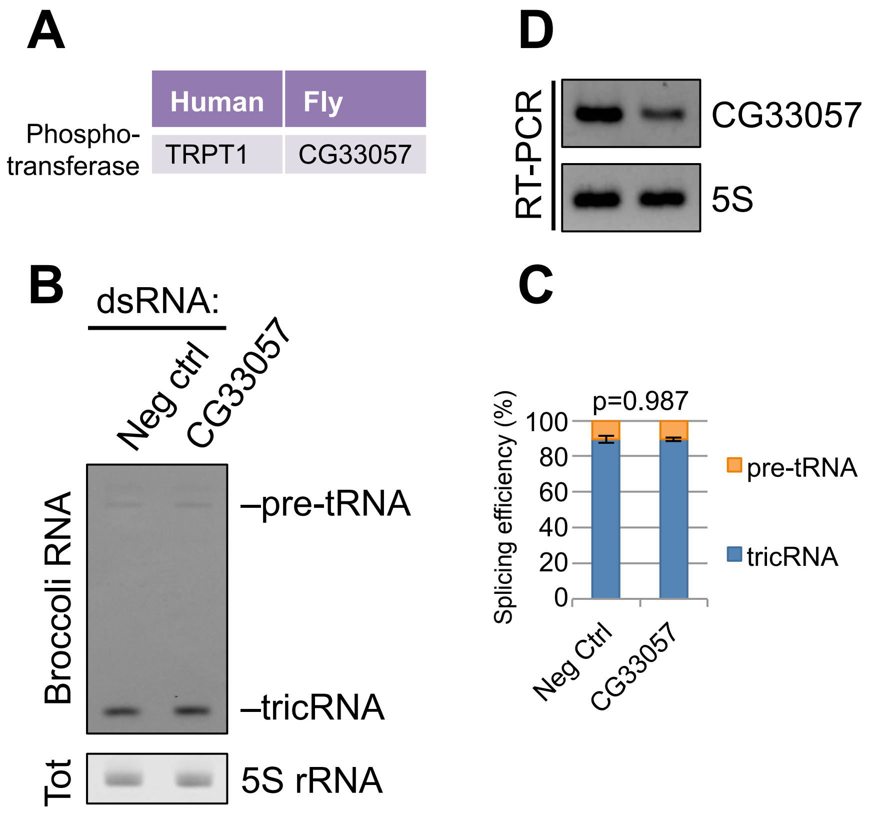


Supplementary figure S4: CG33057 does not participate in reporter tricRNA splicing. (A) *Drosophila* CG33057 is a sequence homolog of human TRPT1. (B) In-gel fluorescence assay of RNA from S2 cells depleted of CG33057. The pre-tRNA and tricRNA bands are identified. The 5S rRNA band from the EtBr-stained gel is shown as a loading control. (C) Quantification of two biological replicates of (B). Error bars denote standard error of the mean. The p-value was calculated using student's t-test. (D) RT-PCR for CG33057 to test knockdown efficiency in S2 cells. RT-PCR for 5S rRNA was used as a control.

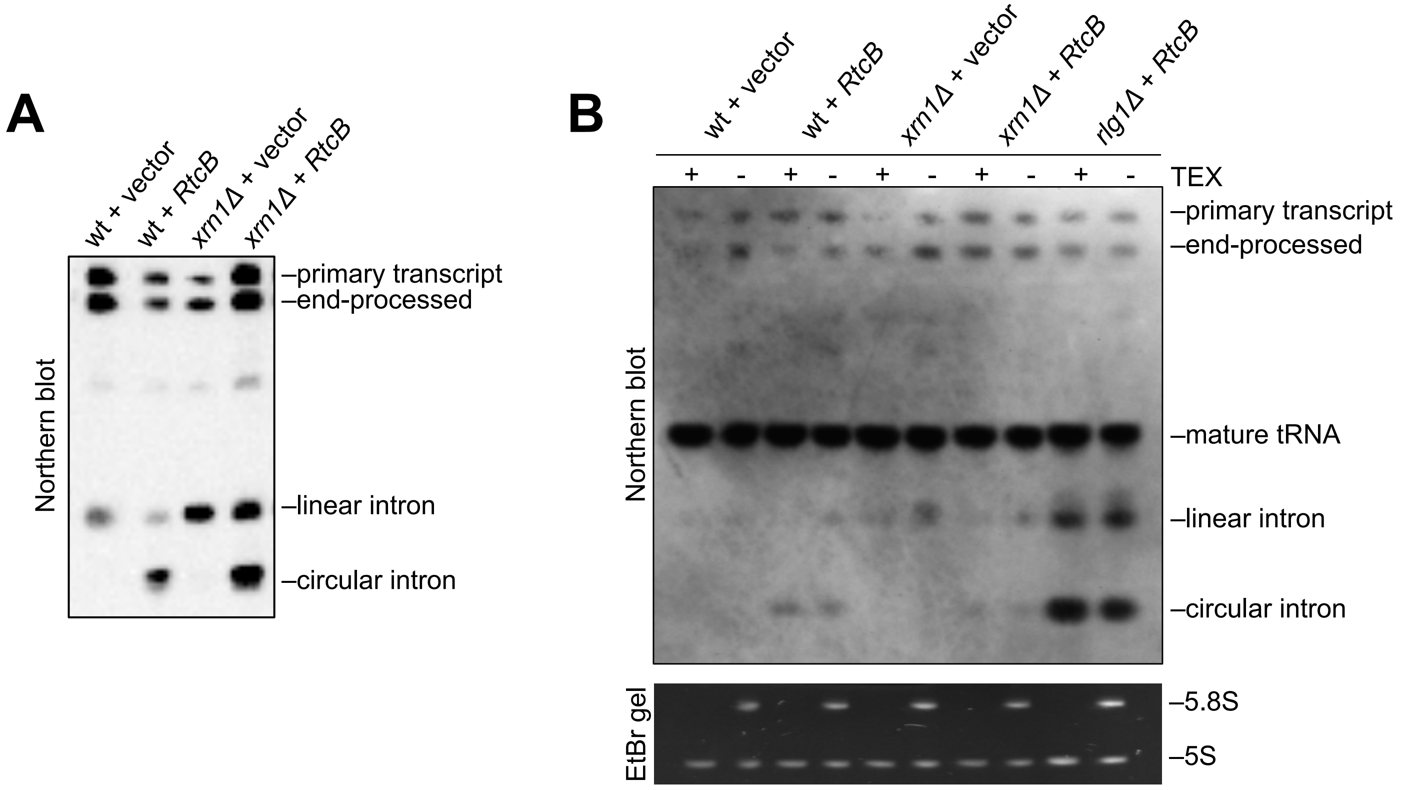


Supplementary Figure S5: Yeast expressing bacterial RtcB generate tricRNAs. (A) Northern blot of RNA from yeast bearing various deletions and enzyme replacements using a probe against tRNA:$Ile_{UAU}$ intron. (B) Northern blot of RNA from yeast bearing various deletions and enzyme replacements using probes for the 5' exon and intron of the tRNA: $Ile_{UAU}$. Top: RNA samples were left untreated (-) or treated (+) with terminator exonuclease (TEX). Bottom: EtBr-stained gel showing 5S and 5.8S as a control.

Yeast lacking the healing and sealing enzyme Rlg1/Trl1 are inviable, but this defect can be complemented by RtcB (11). Notably, there is a clear buildup of circular tRNA introns in the the "RtcB replacement" (*rlg1Δ+RtcB*) strain, along with the linear form. TEX (terminator exonuclease) degrades the majority of linear, unstructured RNAs (e.g. 5.8S rRNA is a good substrate whereas 5S rRNA is refractive to cleavage, see bottom panel of S5B). TEX requires a 5' phosphate for its activity, and because Rlg1/Trl1 also contains the kinase activity that normally phosphorylates the 5' end of the excised intron (6), the linear tRNA introns in the RtcB replacement yeast are not phosphorylated on their 5' ends and are thus poor substrates for TEX.

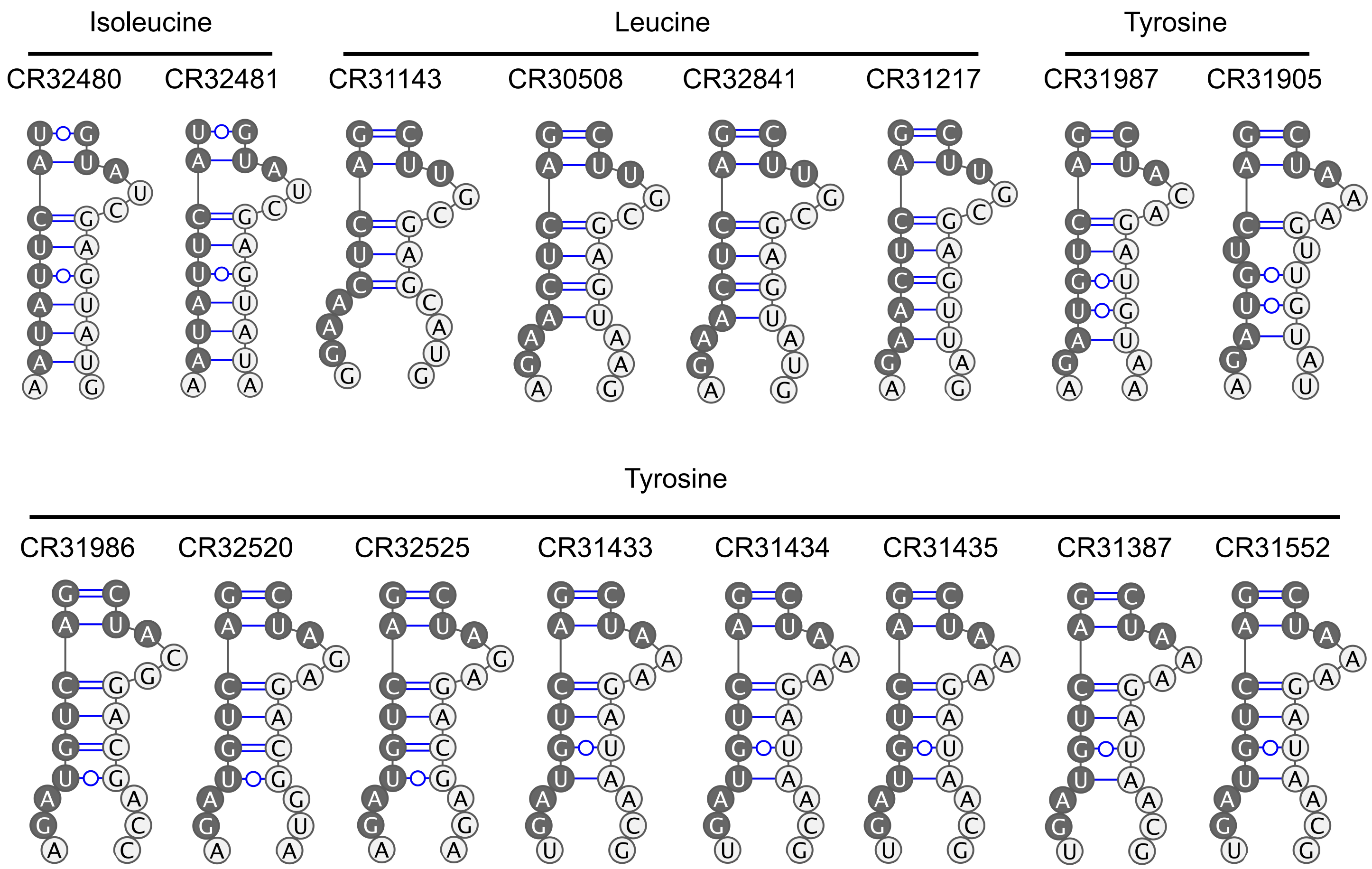


Supplementary Figure S6: Predicted secondary structures of BHB-like motifs for all *Drosophila* intron-containing pre-tRNAs. Sequences of Computed RNA (CR) genes were obtained from FlyBase (http://flybase.org) and the Genomic tRNA Database (http://gtrnadb.ucsc.edu/). CR31905 was discovered after publication of the database. Structures were predicted using Mfold (http://unafold.rna.albany.edu/?q=mfold/RNA-Folding-Form), and structures were drawn using VARNA (http://varna.lri.fr/).

| Purpose | Primer name | Sequence 5'-3' |
|---|---|---|
| Generating dual reporter | Dual rep_F | CCGGCTCGGAGGATATTATTTTGGGTTTTCTTC |
| | Dual rep_R | AATCGAACCGGCGACCTATGGATTTCAAC |
| Northern blot probes | U1 | GAATAATCGCAGAGGTCAACTCAGCCGAGGT |
| | U6 | CTTCTCTGTATCGTTCCAATTTTAGTATATGTTCTGCCGAAGCAAGA |
| | 7SK | TAACCCGTCGTCATCCAGTGGAAGGCAG |
| | Dual reporter probe | TCCGAGCCGGAATCGAACCGG |
| | S.c. tRNA-Ile 5' exon | TATAAGCACGAAGCTCTAACCACTGAGCTACACGAGC |
| | S.c. tRNA-Ile intron | CGTTGCTTTTAAAGGCCTGTTTGAAAGGTCTTTGGCACAGAAACTTCGGAAACCGAATGTTGCTAT |
| Reducing stem length | shortstem5_F | GTATCTGTCGAGTAGAGTGTGGGCTCGCGGTGTTGAAATCCATAGGTCG |
| | shortstem5_R | GAATATCTGGACCCGACCGTCTCGCACTCTACAGTCCACCGC |
| | shortstem6_F | GTATCTGTCGAGTAGAGTGTGGGCTCCGCGGTGTTGAAATCCATAGGT |
| | shortstem6_R | GAATATCTGGACCCGACCGTCTCCGCACTCTACAGTCCACCG |
| | shortstem7_F | GTATCTGTCGAGTAGAGTGTGGGCTCCCGCGGTGTTGAAATCCATAGGT |
| | shortstem7_R | GAATATCTGGACCCGACCGTCTCCCGCACTCTACAGTCCACCG |
| | shortstem8_F | GTATCTGTCGAGTAGAGTGTGGGCTCGCCGCGGTGTTGAAATCCATAGGT |
| | shortstem8_R | GAATATCTGGACCCGACCGTCTCGCCGCACTCTACAGTCCACCG |
| | shortstem9_F | GTATCTGTCGAGTAGAGTGTGGGCTCGGCCGCGGTGTTGAAATCCATAGGT |
| | shortstem9_R | GAATATCTGGACCCGACCGTCTCGGCCGCACTCTACAGTCCACCG |
| | shortstem10_F | GTATCTGTCGAGTAGAGTGTGGGCTCCGGCCGCGGTGTTGAAATCCATAGGT |
| | shortstem10_R | GAATATCTGGACCCGACCGTCTCCGGCCGCACTCTACAGTCCACCG |
| pre-tRNA IVT primers | pre-tRNA_IVT_F | GTATAATACGACTCACTATAGCCTTCGATAGCTCAGTTGGT |
| | pre-tRNA_IVT_R | TCCTTCGAGCCGGATTTGAA |
| Proximal base pair mutations | G-C_F | TATCTGTCGAGTAGAGTGTGGGCTCGTGGCCGCGGTGTTCAAATCCATAGGTCGCTGG |
| | G-C_R | CGAATATCTGGACCCGACCGTCTCGCGGCCGCACTCTACACTCCACCGCTCTACCAACT |
| | A-U_F | TATCTGTCGAGTAGAGTGTGGGCTCGTGGCCGCGGTGTTTAAATCCATAGGTCGCTGG |
| | A-U_R | CGAATATCTGGACCCGACCGTCTCGCGGCCGCACTCTACATTCCACCGCTCTACCAACT |
| | U-A_F | TATCTGTCGAGTAGAGTGTGGGCTCGTGGCCGCGGTGTTAAAATCCATAGGTCGCTGG |
| | U-A_R | CGAATATCTGGACCCGACCGTCTCGCGGCCGCACTCTACAATCCACCGCTCTACCAACT |
| | G-U_F | TATCTGTCGAGTAGAGTGTGGGCTCGTGGCCGCGGTGTTTAAATCCATAGGTCGCTGG |
| | G-U_R | CGAATATCTGGACCCGACCGTCTCGCGGCCGCACTCTACACTCCACCGCTCTACCAACT |
| | U-G_F | TATCTGTCGAGTAGAGTGTGGGCTCGTGGCCGCGGTGTTGAAATCCATAGGTCGCTGG |
| | U-G_R | CGAATATCTGGACCCGACCGTCTCGCGGCCGCACTCTACAATCCACCGCTCTACCAACT |
| | C-C_F | CCGCGGTGTTCAAATCCATAGG |
| | C-C_R | CCACGAGCCCACACTCTA |
| | Dual rep C-C_F | CCGGCTCGGAGGATATTATTTTGGGTTTTCTTC |
| | Dual rep C-C_R | AATCGAACCGGCGACCTATGGATTTGAAC |
| Pairing the helix | 3-UA_F | TGGCCGCGGTATTGAAATCCA |
| | 3-UA_R | CGAGCCCACACTCTACTC |
| | 2-GC_F | GGCCGCGGTGCTGAAATCCAT |
| | 2-GC_R | ACGAGCCCACACTCTACTC |
| | 1-UA_F | GCCGCGGTGTAGAAATCCATA |
| | 1-UA_R | CACGAGCCCACACTCTAC |
| | 3-UA;2-GC_F | TGGCCGCGGTACTGAAATCCATAG |
| | 3-UA;2-GC_R | CGAGCCCACACTCTACTC |
| | 2-GC;1-UA_F | GGCCGCGGTGCAGAAATCCATAG |
| | 2-GC;1-UA_R | ACGAGCCCACACTCTACT |
| | 3-UA;2-GC;1-UA_F | TGGCCGCGGTACAGAAATCCATAG |
| | 3-UA;2-GC;1-UA_R | CGAGCCCACACTCTACTC |
| Unpairing the helix | 4-AA_F | GTGGCCGCGGAGTTGAAATCC |
| | 4-AA_R | GAGCCCACACTCTACTCG |
| | 4-AA;3-UU_F | GTGGCCGCGGATTTGAAATCCATAG |
| | 4-AA;3-UU_R | GAGCCCACACTCTACTCG |
| | 3-UU;2-GG_F | TGGCCGCGGTTGTGAAATCCATAG |
| | 3-UU;2-GG_R | CGAGCCCACACTCTACTC |
| | 4-AA;3-UU;2-GG_F | GTGGCCGCGGATGTGAAATCCATAGGTC |
| | 4-AA;3-UU;2-GG_R | GAGCCCACACTCTACTCG |
| | 4-AA;3-UU;2-GG;1-UA_F | GTGGCCGCGGATGAGAAATCCATAGG |
| | 4-AA;3-UU;2-GG;1-UA_R | GAGCCCACACTCTACTCG |
| Human tricRNA reporter *cis* element mutations | C-C_F | TGTTCAAATCCATAGGTCGCTGGTTCAAATCCGGC |
| | C-C_R | CCGCGGCCACGAGCC |
| | 4-AA_F | AGTTGAAATCCATAGGTCGCTGGTTCAAATCCGGC |
| | 4-AA_R | CCGCGGCCACGAGCC |
| | 3-UA_F | TATTGAAATCCATAGGTCGCTGGTTCAAATCCGGC |
| | 3-UA_R | CCGCGGCCACGAGCC |
| | 2-GC_F | TGCTGAAATCCATAGGTCGCTGGTTCAAATCCGGC |
| | 2-GC_R | CCGCGGCCACGAGCC |
| | 3-UA;2-GC_F | TACTGAAATCCATAGGTCGCTGGTTCAAATCCGGC |
| | 3-UA;2-GC_R | CCGCGGCCACGAGCC |

Supplementary Table 1: List of oligonucleotides used in this study

| Purpose | Primer name | Sequence 5'-3' |
|---|---|---|
| Making PCR products for *in vitro* transcription of *Drosophila* processing factors | TSEN2_F | TAATACGACTCACTATAGGGGGTATTAAGTTCGGCGGTGATTTTCG |
| | TSEN2_R | TAATACGACTCACTATAGGGCTTCTTAGGCGGTTGGACAGTC |
| | TSEN15_F | TAATACGACTCACTATAGGGGCTGTTGTGAATCTGGCACAGA |
| | TSEN15_R | TAATACGACTCACTATAGGGATCATCGGGGCTTACAGTTGTG |
| | TSEN34_F | TAATACGACTCACTATAGGGCGAACACAAAAGGCTGGAGTC |
| | TSEN34_R | TAATACGACTCACTATAGGGACTCCTCAAGTTTCTCGGCA |
| | TSEN54_F | TAATACGACTCACTATAGGCGTTTACAACTGGAGTACTGTGGTTT |
| | TSEN54_R | TAATACGACTCACTATAGGGCACCAAATCGAACTTTTCAAAGATC |
| | Ddx1_F | TAATACGACTCACTATAGGGCGCCTAACGCCCCTCA |
| | Ddx1_R | TAATACGACTCACTATAGGGCGGGTCGACTAGACAGACGA |
| | Archease_F | TAATACGACTCACTATAGGGCCACGGATGGGGATCGT |
| | Archease_R | TAATACGACTCACTATAGGGAATGTCAATTATCACGAACACCTCGTAGT |
| | RtcB_F | TAATACGACTCACTATAGGACGCCGAGATCCAGGTGG |
| | RtcB_R | TAATACGACTCACTATAGGAACGCTTGACGGGTGAGGAATG |
| | CG33057_F | TAATACGACTCACTATAGGGCGTTCCCGATTTGCAGAAGCA |
| | CG33057_R | TAATACGACTCACTATAGGGTCCGCCAGCACCTTTTCC |
| | Zucchini_F | TAATACGACTCACTATAGGGAGCAAGCGAGAGAAGGCAAG |
| | Zucchini_R | TAATACGACTCACTATAGGGCCAAGAGCCGTCCAGTTTACG |
| | Smg6_F | TAATACGACTCACTATAGGGATTGCTGGGCTGAGCTAACAA |
| | Smg6_R | TAATACGACTCACTATAGGGTGTCCACGAACTTGAGAATGTCC |
| | Dis3_F | TAATACGACTCACTATAGGGCTTGCCGAAAATGCCCTGGACAATTA |
| | Dis3_R | TAATACGACTCACTATAGGGATCGCTCAACACCGCCAACC |
| | Clipper_F | TAATACGACTCACTATAGGGGGACCGGACGATCGTGT |
| | Clipper_R | TAATACGACTCACTATAGGGCCGGAGTGTCCAGAGTAGC |
| | Neg ctrl_F | TAATACGACTCACTATAGGGTTCAACATCGTGGCCGTGGC |
| | Neg ctrl_R | TAATACGACTCACTATAGGGGTTGGCAAGCCCTTTGAGGCA |
| *Drosophila* processing factor RT-PCR primers | TSEN2_F | TGGTTGTTTTGGCAAGGGAAGCA |
| | TSEN2_R | ACCTCCACAAACATGCAGAAGTCC |
| | TSEN15_F | GATTTGACCGCCGCTTTGGG |
| | TSEN15_R | GCCTTTGTGTGCAGCACAGG |
| | TSEN34_F | GGTACTGGCTTCGTTTTCAACGTGG |
| | TSEN34_R | GTTTCAAATTTACTGAGCTCCACGGGC |
| | TSEN54_F | GGAGCTTAAACGAGCGCAGGAGTA |
| | TSEN54_R | GCTTTCCCTGCTCACTGTATCCGAA |
| | Ddx1_F | CAGACGAATTGGACTGGACCCTG |
| | Ddx1_R | GGTCTCCCACACGATTTGCAGAA |
| | Arch_F | CTGGATCACACGGCGGATGTTC |
| | Arch_R | CTCTCCAGATCATCTCCATGCGC |
| | RtcB_F | CTTCCCGCCACACCATCC |
| | RtcB_R | CACGACCCGCTCCGTG |
| | CG33057_F | GGAATCACGATCCGTGCTGATGG |
| | CG33057_R | CGAAGAGTATATCGTTGCTTGGCATCC |
| | Zucchini_F | CCTCAGAGGTGATTTGGAAGCTGG |
| | Zucchini_R | CGACACTTGTGGTTTCTGGGATCC |
| | Smg6_F | CGAGCACAACAAACCCAAAACGTCA |
| | Smg6_R | GCAAAGTCTTCTTCTGGGAGACACTG |
| | Dis3_F | CGAGCACCACAAGGAAACCTATGC |
| | Dis3_R | CTCTTTAGAGGATTGCGAGGCCTG |
| | Clipper_F | CATCACCAGGAACGGACAGGAG |
| | Clipper_R | GTAGCACTCGGGCATCTTGGT |
| | 5S_F | AACAACACGCGGTGTTCCCAAGC |
| | 5S_R | GCCAACGACCATACCACGCTGAA |
| Gateway cloning *Drosophila* endonucleases | Dis3_N_F | GGGGACAAGTTTGTACAAAAAAGCAGGCTTCCAAACTTTACGCGAATTTAC |
| | Dis3_N_R | GGGGACCACTTTGTACAAGAAAGCTGGGTNTTACTTCTTTTTCTTATCCTTCT |
| | Dis3_C_F | GGGGACAAGTTGGTACAAAAAAGCAGGCTATGCAAACTTTACGCGAA |
| | Dis3_C_R | GGGGACCACTTTGTACAAGAAAGCTGGGTCCTTCTTTTTCTTATCCTTCTTTGTC |
| | Clp_N_F | GGGGACAAGTTTGTACAAAAAAGCAGGCTTCGACATCCTTTTGGCCAAC |
| | Clp_N_R | GGGGACCACTTTGTACAAGAAAGCTGGGTNCTACTTATGACTATGTTGGTTAGAG |
| | Clp_C_F | GGGGACAAGTTGGTACAAAAAAGCAGGCTATGGACATCCTTTTGGCC |
| | Clp_C_R | GGGGACCACTTTGTACAAGAAAGCTGGGTCCTTATGACTATGTTGGTTAGAGAGG |
| Yeast RT-PCR primers | S.c. Ile_F | TGCTTTTAAAGGCCTGTTTGAAAGG |
| | S.c. Ile_R | GCAACATTCGGTTTCCGAAGTTTCT |

Supplementary Table 1 continued: List of oligonucleotides used in this study
